# A p53-ΔNp73 signaling axis drives selective motor neuron degeneration in spinal muscular atrophy

**DOI:** 10.64898/2026.06.20.733537

**Authors:** Maria J Carlini, Xuemei Hu, Leonie Sowoidnich, Archana Yadav, Jason Q Garcia, Maria Savin, Charlotte J Sumner, Vilas Menon, Christian M Simon, Livio Pellizzoni

## Abstract

Selective neuronal vulnerability is a hallmark of many neurodegenerative diseases, yet how ubiquitous genetic insults cause highly selective neuronal loss remains poorly understood. In spinal muscular atrophy (SMA), reduced SMN levels trigger degeneration of specific motor neuron pools. Although non-apoptotic, p53-mediated death pathways have been implicated, p53 is expressed in both vulnerable and resistant neurons, leaving the downstream determinants of selective vulnerability unresolved. Here, we identify a p53–ΔNp73 signaling axis as a previously unrecognized execution pathway driving motor neuron degeneration. Using differential transcriptional profiling of SMA motor neurons following pharmacological modulation of p53 activity, we uncover p73 as a critical downstream mediator of neuronal death. Notably, SMN deficiency induces cell-autonomous, p53-dependent expression of the ΔNp73 isoform selectively in vulnerable, but not resistant, motor neurons. ΔNp73 induction precisely parallels the spatial and temporal pattern of degeneration in mouse models and is also detected in motor neurons from SMA patients. Strikingly, despite its established role as a pro-survival antagonist of p53, depletion of ΔNp73 improves motor neuron survival and partially preserves neuromuscular junction integrity in SMA mice. These findings reveal a context-dependent, isoform-specific functional switch in p53 family signaling that redirects a canonical survival factor into a driver of neurodegeneration, identifying a novel molecular mechanism underlying selective neuronal vulnerability in SMA and a potential therapeutic target for neuroprotection.

## Introduction

Selective neuronal vulnerability is a defining but unresolved feature of many neurodegenerative diseases^1,2^. Across diverse genetic etiologies, degeneration often affects only specific neuronal subtypes despite widespread molecular insults, suggesting that cell-intrinsic programs dictate selective susceptibility to otherwise ubiquitous disease triggers. Defining these programs remains a central challenge in neuroscience, as it is essential for understanding disease progression and developing targeted neuroprotective strategies. However, the molecular logic linking broadly expressed disease-causing pathways to highly selective neuronal loss remains poorly understood. Here, we investigated this question in spinal muscular atrophy (SMA), a prototypical model in which a ubiquitous genetic defect leads to strikingly selective neurodegeneration^3,4^.

SMA is a neuromuscular disorder caused by homozygous deletion or mutation of the *survival motor neuron 1* (*SMN1*) gene^5^, with disease severity modified by copy number of the paralogous *SMN2* gene^3,4^. Unlike *SMN1*, the *SMN2* gene predominantly produces a truncated and unstable protein due to exclusion of exon 7^6,7^, thereby providing only partial functional compensation. SMN functions as the core component of a multi-protein complex required for the assembly of small nuclear ribonucleoproteins (snRNPs), essential for pre-mRNA splicing and histone mRNA 3’-end processing^3^. In addition, SMN has been implicated in other aspects of RNA metabolism, including mRNA transport and translation^8,9^. Despite these ubiquitous cellular functions, a striking feature of SMA is the selective degeneration of spinal motor neurons. This vulnerability is not uniform, however, as motor neurons innervating proximal and axial muscles are particularly susceptible, whereas those controlling distal muscles are comparatively resistant^10,11^. This selective degeneration is faithfully recapitulated in the SMNΔ7 mouse model^12^, which has been instrumental in defining disease mechanisms and testing therapeutic strategies.

A central pathway implicated in SMA motor neuron degeneration is activation of the transcription factor p53^13^. Although p53 is classically associated with genome surveillance and tumor suppression^14^, it has also been increasingly linked to neurodegenerative disease^15^, including the adult onset motor neuron disease amyotrophic lateral sclerosis (ALS)^16–19^. In SMNΔ7 mice, vulnerable motor neurons exhibit early nuclear accumulation of p53 and phosphorylation at serine 18^13^, both of which precede overt cell death. In contrast, resistant motor neurons display delayed p53 accumulation and lack serine 18 phosphorylation^13,20,21^. Mechanistically, SMN deficiency disrupts snRNP assembly and leads to specific splicing defects that converge on p53 regulation. Dysregulated alternative splicing of Mdm2 and Mdm4 promotes p53 stabilization and nuclear accumulation^22^, while impaired minor intron splicing of Stasimon enhances p38 mitogen-activated protein kinase (MAPK) signaling^23^, which contributes to p53 serine 18 phosphorylation. As inhibition of either p53 or p38 MAPK provides neuroprotection^13,20,23,24^, these signaling pathways converge to drive selective degeneration of vulnerable SMA motor neurons.

Despite this progress, the downstream mechanisms by which p53 executes motor neuron death in SMA are unknown. Surprisingly, degenerating motor neurons do not exhibit canonical features of p53-mediated apoptosis, including caspase activation and DNA fragmentation^13^. This suggests that p53 may trigger a non-canonical form of neuronal death in the context of SMA. Thus, while upstream mechanisms leading to p53 activation are increasingly well defined, the downstream effectors that execute degeneration—and potentially determine selective vulnerability—remain unknown.

This gap is particularly important in the current clinical landscape of disease-modifying therapies that restore SMN expression, including viral-mediated gene delivery^25^, antisense oligonucleotides^26,27^, and small-molecule splicing modifiers^28,29^. Although these interventions have transformed SMA treatment, they do not fully prevent motor neuron loss or restore neuromuscular function, especially when administered after symptom onset^30–32^. These limitations suggest that SMN deficiency initiates pathogenic programs that become partially independent of SMN restoration, underscoring the need to identify downstream effectors of neurodegeneration as targets for combinatorial therapeutic strategies.

Here, we addressed this question by identifying p53-dependent transcriptional targets that execute motor neuron death in SMA. Using differential transcriptional profiling of SMA motor neurons under conditions of pharmacological inhibition of p53 and p38 MAPK signaling, we identified p73 – a p53 family transcription factor – as a selective marker of vulnerable motor neurons and a key downstream effector of degeneration. Strikingly, SMN deficiency induced p53-dependent expression of the ΔNp73 isoform, which lacks the amino terminal transactivation domain, specifically in vulnerable SMA motor neurons. Functional inhibition of ΔNp73 significantly improved motor neuron survival and neuromuscular junction innervation in SMA mice. Although ΔNp73 has previously been characterized as a factor that antagonizes p53 activity through dominant negative mechanisms^33,34^, our findings reveal a context-dependent switch in which ΔNp73 functions downstream of p53 to promote neuronal death. These results define a novel p53–ΔNp73 signaling axis as a critical execution pathway in SMA and provide new insight into the mechanisms underlying selective motor neuron vulnerability, with potential implications for other neurological disorders involving p53.

## Results

### Identification of candidate effectors of p53-dependent motor neuron death in SMA mice

To identify the downstream executioners of p53-dependent motor neuron death, we performed differential profiling of gene expression in motor neurons isolated from SMNΔ7 SMA mice treated with selective inhibitors of p53 (pifithrin-α, PFTα) and p38α MAPK (MW150) relative to untreated WT and SMA mice. We also included motor neurons isolated from SMA mice treated with the SMN-inducing compound SMN-C3 - an analog of Risdiplam that is clinically approved for SMA treatment^35^ – as an additional experimental group for SMN-dependent correction of transcriptome changes. Newborn SMA mice were injected intraperitoneally (IP) with PFTα (2.2 mg/kg), MW150 (5mg/kg) or SMN-C3 (3mg/kg) once daily from birth to P6 (Figure 1A), a time of robust p53 activation and ongoing motor neuron degeneration in the SMNΔ7 model^13^. To specifically identify L1-L3 motor neurons directly innervating a disease-vulnerable muscle for laser capture microdissection (LCM), we used retrograde tracing with fluorescently labeled Cholera Toxin b (CTB) injected in the quadratus lumborum (QL) and iliopsoas (IL) muscles at P2 followed by spinal cord dissection at P6 (Figure 1A) as in our previous studies^22,36^. We then performed isolation of motor neurons by LCM, total RNA purification, and transcriptome-wide analysis of gene expression by RNA-seq. Gene expression analysis identified 184 differentially expressed genes (50 down-regulated and 134 up-regulated) between WT and SMA motor neurons (Figure 1B and Supplementary Table S1). Pathway analysis identified activation of p53 signaling as the top enriched biological process in this dataset (Figure 1F), which is consistent with the results of our earlier studies^13,22^. Next, we asked which gene changes were significantly corrected in motor neurons by treatment of SMA mice with PFTα and/or MW150 relative to untreated SMA motor neurons (Figure 1C-E and Supplementary Table S1), which identified 18 genes (Figure 1G). Further refinement by selection of those genes that were also corrected by treatment with SMN-C3 yielded a narrower group of 8 genes (Figure 1G and H). These genes (*Atp11c*, *Atp2b4*, *B3gnt8*, *Dzip3*, *Emp3, Sytl1*, *Trp73 and Zfp119b*) are dysregulated in SMA motor neurons and significantly corrected by either SMN induction or p53 inhibition (Figure 1B-D). There is also modulation of these genes by p38 MAPK inhibition with MW150 (Figure 1E and H), although the correction did not reach statistical significance. The changes in the expression of 6 out of the 8 genes (a probe for *Zfp119b* could not be designed, and that for *B3gnt8* did not yield any signal) were validated by RNAscope analysis of their corresponding mRNAs in lumbar L1 motor neurons from WT and SMA mice at P6 (Figure 2A and 2B and Supplementary Figure S1). Of note, SMN restoration by SMN-C3 corrected a much larger proportion of SMA-related changes (74 genes) relative to p53 or p38 MAPK inhibition as expected, but also elicited a greater number of unrelated gene changes (481 genes), indicating a significant off target impact of this drug on the motor neuron transcriptome (Figure 1G) that is consistent with findings in cultured cells^37^. In sum, this analysis identified a set of genes as candidates to mediate the p53-dependent death of SMA motor neurons induced by SMN deficiency for further functional analysis.

**Figure 1.**
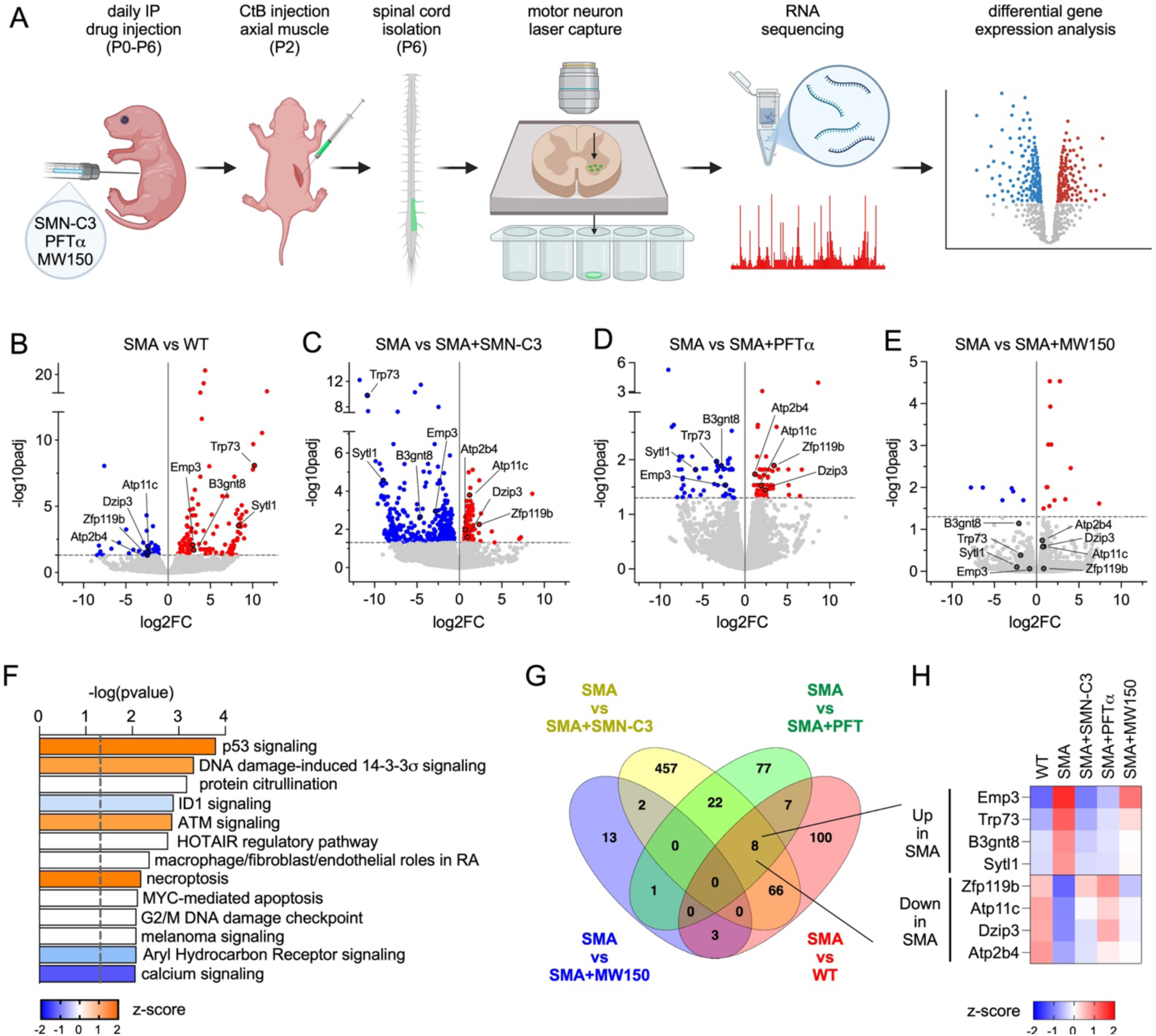
Identification of candidate effectors of p53-dependent motor neuron death in SMA mice. (**A**) Schematic of the experimental design to identify candidate effectors of SMA motor neuron death. See text for details. (**B-E**) Volcano plots of the indicated two-way comparisons of transcript-level changes in motor neurons isolated from P6 WT and SMA mice that were either untreated or treated daily with the indicated drugs (SMN-C3, PFTα, or MW150) from birth. For each comparison, significant gene changes (q<0.05) are indicated in blue (downregulated) and red (upregulated). The 8 candidate effector genes whose expression is dysregulated in SMA motor neurons and corrected upon SMN induction or p53 inhibition are shown. See also Supplementary Table S1. (**F**) Ingenuity pathway analysis of dysregulated genes in SMA motor neurons relative to WT as controls. The z-score color coding visualizes predictions for the pathway(s) to be activated (z>0) or inhibited (z<0) and the dashed line indicates the significance threshold. (**G**) Venn diagram of significant transcript-level changes (q<0.05) identified in the two-way comparisons of the indicated experimental groups. (**H**) Heatmap of the relative expression of the 8 candidate effector genes across experimental groups.

**Figure 2.**
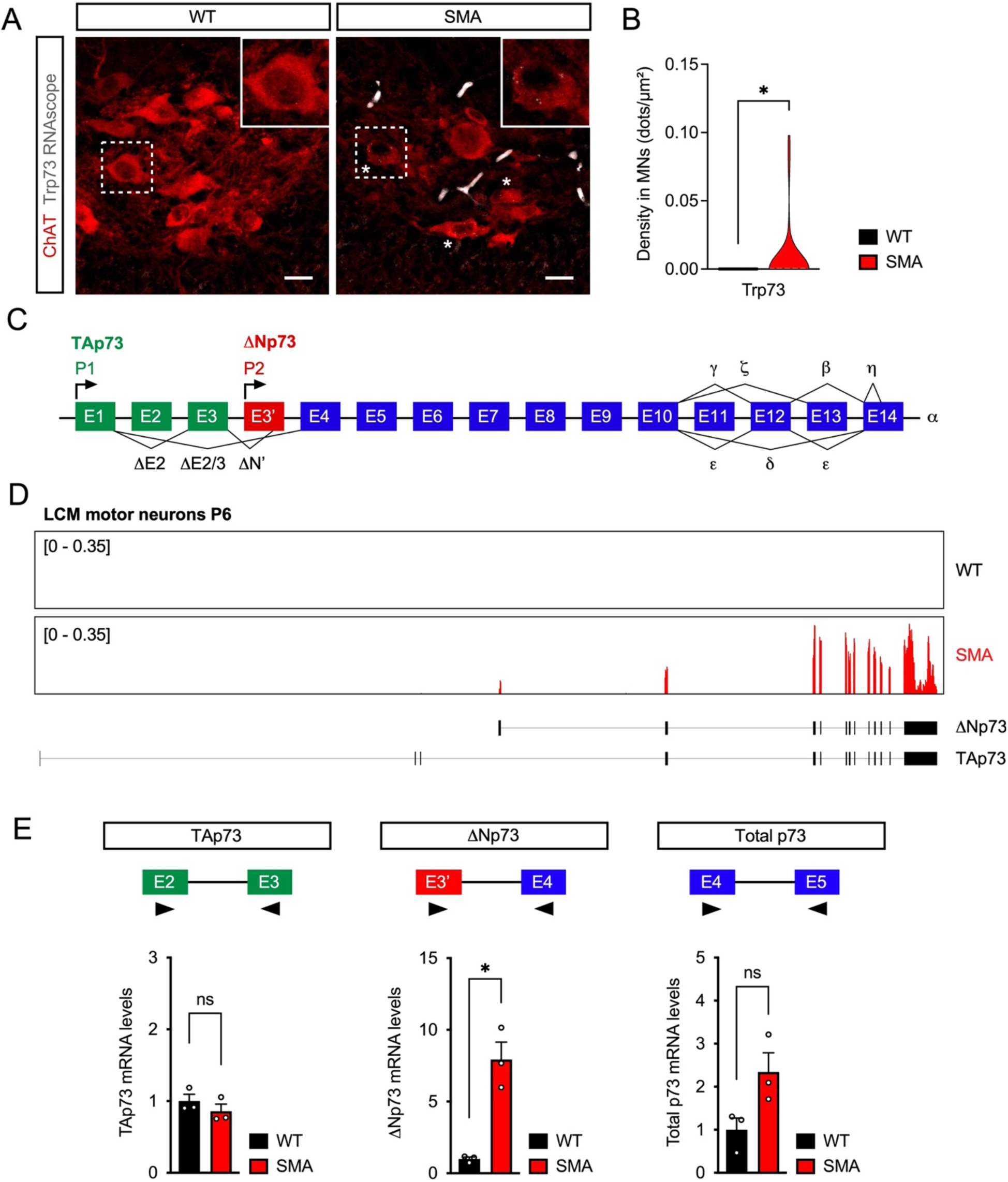
SMN deficiency induces ΔNp73 mRNA in motor neurons of SMA mice. (**A**) ChAT immunostaining and RNAscope detection of p73 mRNAs in L1 spinal cord sections from WT and SMA mice at P6. Scale bars = 20 µm. Asterisks mark the subset of SMA motor neurons with detectable p73 mRNA foci. Magnified views of the boxed areas are shown in the insets. (**B**) Violin plot of the number of p73 mRNA foci normalized to the soma area (dots/μm^2^) from WT (n=30) and SMA (n=30) motor neurons. Mann-Whitney test. *P < 0.05. (**C**) Schematic of the exon-intron organization of the mouse *Trp73* gene showing the transcription initiation sites for TAp73 (P1) and ΔNp73 (P2) mRNAs. Green and red exons are specific to TAp73 and ΔNp73 mRNAs, respectively. Blue exons are common to both TAp73 and ΔNp73 mRNAs and alternative splicing events yielding distinct transcript isoforms are also shown. (**D**) Representative RNA-seq tracks for reads mapped to the *Trp73* locus from motor neurons of WT and SMA mice at P6. Reference transcript structures of TAp73 and ΔNp73 are shown. (**E**) RT-qPCR analysis of TAp73, ΔNp73 and total p73 mRNAs in P11 spinal cords from WT and SMA mice. The graph shows mean, SEM, and individual values normalized to WT as a control (n=3 mice). Two-tailed unpaired Student’s t-test for TAp73 and Total p73 mRNAs, Welch’s test for ΔNp73 mRNA. *P < 0.05; ns, not significant. The schematics show the location of the PCR primers.

### Isoform-specific p73 expression selectively marks vulnerable motor neurons in SMA mice

The genes identified in our differential screening may represent direct transcriptional targets of p53 or indirect downstream changes of functional networks regulated by p53 as well as changes of motor neuron pathology. To begin addressing these possibilities, we prioritized our follow-up analysis on the characterization of the *Trp73* gene as a candidate marker and executioner of motor neuron death in SMA. This choice was guided by the fact that *Trp7*3 is a transcriptional target of p53 with established functions in regulating developmental processes and programmed cell death pathways^34,38,39^.

The *Trp73* gene generates two main protein isoforms - TAp73 and ΔNp73^34^ - that differ at the amino terminus due to usage of distinct transcription initiation sites (Figure 2C) and harbor or lack the transactivation domain (Supplementary Figure S2A), respectively. Moreover, alternative splicing provides further diversification of each main isoform (Figure 2C)^34,40^. Importantly, both the analysis using RNAscope probes and the mapping of reads from RNA sequencing demonstrated that WT motor neurons do not express *Trp73* above detection levels, while low levels of p73 mRNA are specifically detected at least in a subset of SMA motor neurons (Figure 2A-D). Interestingly, only transcripts corresponding to the ΔNp73 isoform accumulate in SMA motor neurons (Figure 2D). To extend this observation, we designed primer sets to detect specifically either TAp73 or ΔNp73 transcripts as well as total p73 mRNAs and performed RT-qPCR analysis of spinal cord RNA isolated from WT and SMA mice at P11 (Figure 2E). This analysis showed a robust and specific induction of ΔNp73 but not TAp73 mRNAs in the SMA spinal cord. Total p73 mRNA levels showed a non-significant increase that was smaller in magnitude than the rise in ΔNp73 mRNA. This lesser impact likely occurs because TAp73 mRNA is the prominent isoform expressed in spinal cord ependymal cells under normal conditions^38,41^. Together, these results indicate that SMN deficiency induces the selective upregulation of ΔNp73 mRNA in SMA motor neurons.

Next, we investigated the expression of the p73 protein and whether it correlated with the selective vulnerability of motor neurons in SMA. We first validated a commercial antibody in human cell lines transfected with HA-tagged mouse TAp73 or ΔNp73 by Western blotting and immunofluorescence analysis (Supplementary Figure S2B and S2C). This antibody recognizes both p73 isoforms, and we were unable to identify an isoform-specific antibody. We then performed immunohistochemistry experiments in spinal cord sections isolated from WT and SMA mice at early (P1), mid (P6), and late (P11) symptomatic stages of the disease in the SMNΔ7 model. We analyzed sections from lumbar L1 segments containing vulnerable motor neurons that degenerate in SMA as well as lumbar L5 segments containing both medial motor column (MMC) and lateral motor column (LMC) motor neurons that are, respectively vulnerable and resistant to death in SMA^12^. We performed immunostaining with antibodies against ChAT to identify motor neurons and p73, followed by confocal microscopy and quantification (Figure 3). Consistent with the analysis of mRNA expression (Figure 2A and 2D, and Supplementary Table S1), we found no evidence of p73 expression in either L1 or L5 motor neurons from WT mice at any of the time points analyzed (Figure 3A-B). In contrast, we identified nuclear accumulation of p73 in vulnerable L1 and L5 MMC motor neurons but not in resistant L5 LMC motor neurons from SMA mice as early as P1 (Figure 3A-B). The percentage of p73^+^ motor neurons in vulnerable SMA motor neuron pools progressively increased from birth to later stages of the disease (Figure 3C), while p73 was only seldom found in few resistant motor neurons even at disease end stage. Taken together, these findings identify ΔNp73 expression as a novel, specific marker of vulnerable SMA motor neurons in a severe mouse model of the disease.

**Figure 3.**
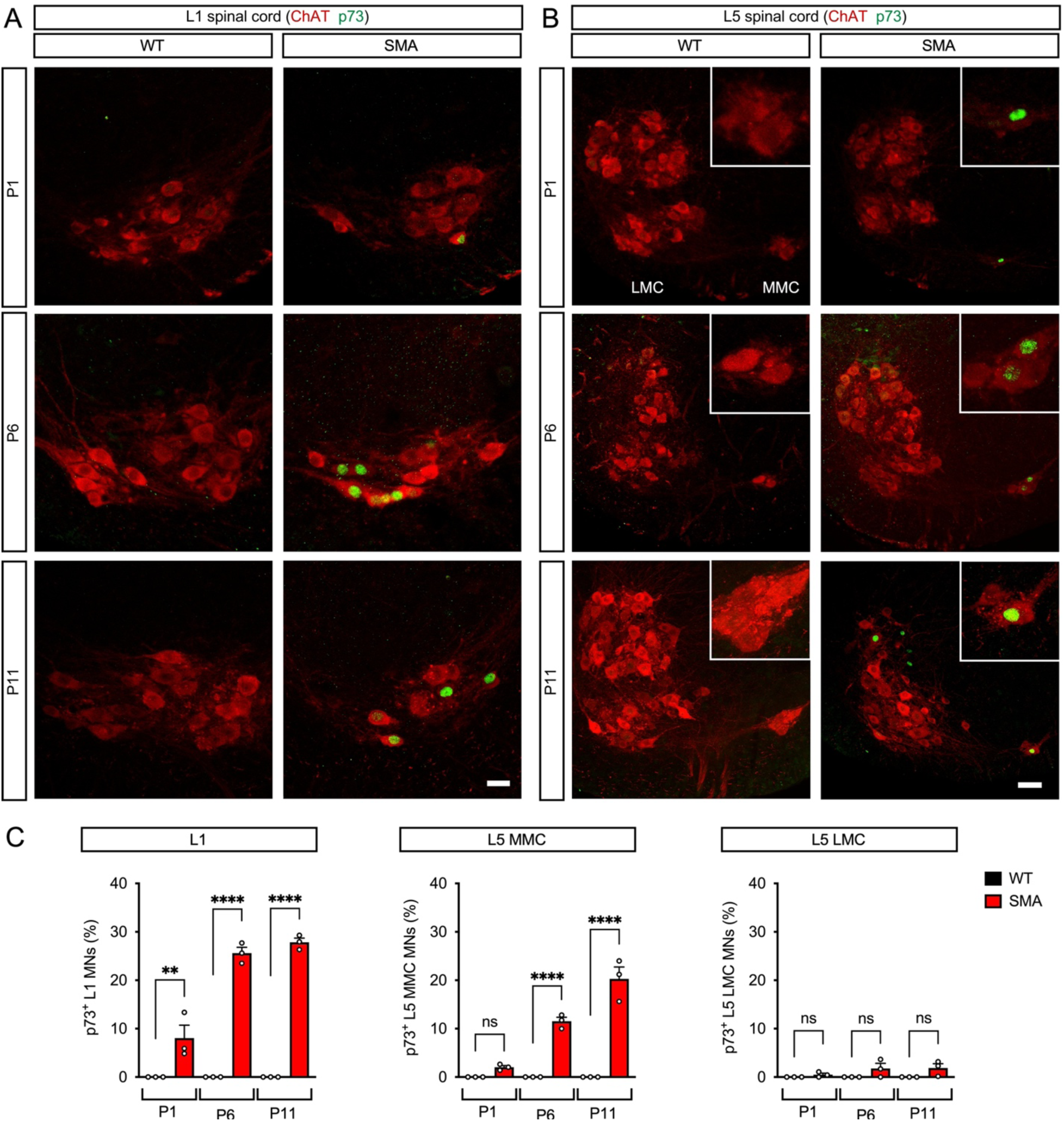
p73 upregulation selectively marks vulnerable motor neurons in SMA mice. (**A**) ChAT and p73 immunostaining of L1 spinal cords isolated from WT and SMA mice at P1, P6, and P11. Scale bar = 25 µm. (**B**) ChAT and p73 immunostaining of L5 spinal cords isolated from WT and SMA mice at P1, P6, and P11. L5 LMC and MMC motor neuron pools are indicated, and magnified views of L5 MMC motor neurons are shown in the insets. Scale bar = 50 µm. (**C**) Percentage of p73^+^ L1, L5 LMC, and L5 MMC motor neurons in WT and SMA mice at P1, P6, and P11. The graphs show mean, SEM, and individual values from WT (n=3) and SMA (n=3) mice. Two-way ANOVA with Sidak’s *post hoc* test. ****P < 0.0001; **P < 0.01; ns, not significant.

### p73 accumulates in p53-expressing motor neurons of a mouse model and type 1 SMA patients

We previously demonstrated that p53 expression in SMA mice is initially restricted to vulnerable motor neuron pools and later extends to a much larger population of spinal neurons^13^, including resistant motor neuron pools and dorsal interneurons. Accordingly, p53 immunostaining of L1 spinal cords shows broad p53 expression beyond motor neurons in the spinal cord of SMA mice at P11 (Figure 4A). In striking contrast, p73 expression remains essentially confined to SMA motor neurons even at this late stage of the disease (Figure 4B). Ependymal cells in the central canal express p73 in both WT and SMA mice as expected^42^. We then investigated the overlap of p53 and p73 expression by co-immunostaining in the spinal cord of SMA mice at P11 and found that all p73^+^ L1 motor neurons also expressed p53 (Figure 4C), while p73 was present in about half of the p53^+^ motor neurons (59.3±9.7% from n=5 mice). Together, these findings show that p73 induction is restricted to a subset of p53-expressing vulnerable SMA motor neurons.

**Figure 4.**
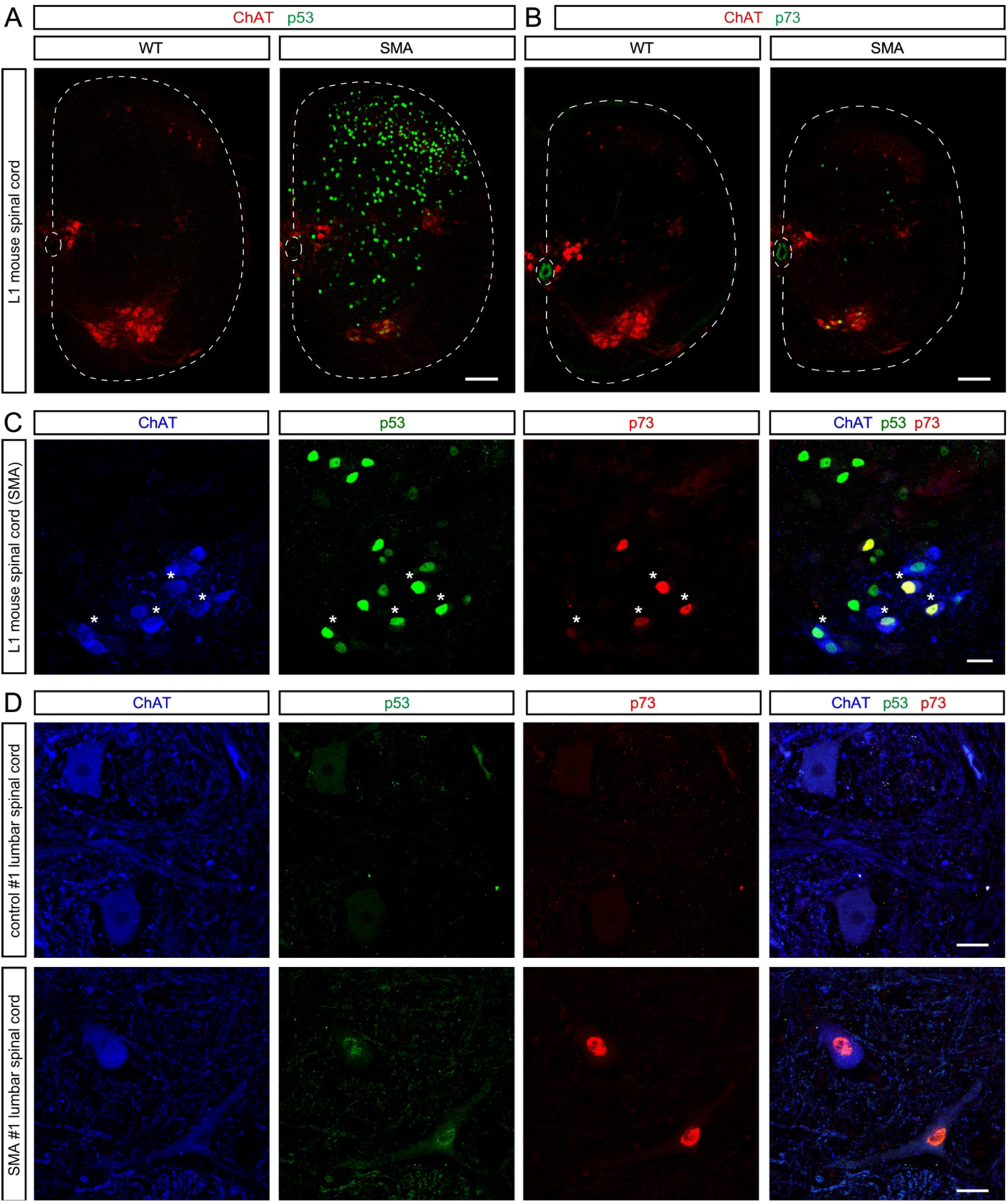
p73 accumulates in p53-expressing motor neurons of a mouse model and type 1 SMA patients. (**A**) ChAT and p53 immunostaining of L1 spinal cords isolated from WT and SMA mice at P11. Scale bar = 100 µm. (**B**) ChAT and p73 immunostaining of L1 spinal cords isolated from WT and SMA mice at P11. Scale bar = 100 µm. (**C**) ChAT, p53 and p73 immunostaining of a lumbar L1 spinal cord isolated from an SMA mouse at P11. Scale bar = 25 µm. Asterisks mark overlapping p53 and p73 expression in motor neurons. (**D**) ChAT, p53 and p73 immunostaining of lumbar spinal cords isolated postmortem from control and severe SMA type 1 individuals. Scale bars = 25 µm. See Supplementary Table S2 for further information about the human samples.

Next, we sought to address the clinical relevance of our findings by investigating the expression of p73 in *postmortem* spinal cords from controls and severe type 1 SMA patients (Supplementary Table S2). Immunohistochemistry and confocal microscopy analysis showed that ChAT^+^ motor neurons from lumbar and thoracic spinal cords of unaffected individuals did not express p73. In contrast, nuclear accumulation of p73 was found in both lumbar and thoracic motor neurons from type I SMA patients (Figure 4D and Supplementary Figure S3). Similar to the findings in the SMNΔ7 mouse model, nuclear accumulation of p53 was also observed in motor neurons from SMA but not control human spinal cords (Figure 4D). Together, these experiments indicate that p73 and p53 induction in motor neurons is a conserved feature in a mouse model of SMA and individuals affected by the most severe form of the disease.

### SMN deficiency drives cell autonomous p73 expression in motor neurons

To determine whether p73 expression originates from cell autonomous effects of SMN deficiency in motor neurons, we investigated whether selective depletion of SMN in motor neurons is necessary and sufficient to induce both p53 and p73 *in vivo*. To do so, we used mice homozygous for a conditional *Smn* allele (*Smn^F^*^7^), which is functionally inactivated upon Cre recombination^43^, also harboring two copies of the human *SMN2* gene for constitutive expression of low SMN levels akin to SMA. These *Smn^F7/F^; SMN2^+/+^* mice were used as controls (no Cre) and crossed with ChAT-Cre mice to obtain experimental mice (*Smn^F7/F7^; SMN2^+/+^; ChAT-Cre^+/-^*) with selective inactivation of mouse *Smn* in cholinergic neurons. To validate that these mice recapitulated motor neuron cell autonomous features of SMA^44,45^, we first analyzed the number of L1 motor neurons and the innervation of neuromuscular junctions (NMJs) in the QL, an axial muscle innervated by this disease-vulnerable motor neuron pool^13,46^. We found a significant loss of L1 motor neurons (>30%) as well as robust NMJ denervation of the QL in ChAT-Cre mice relative to no-Cre controls at P13 (Figure 5A-D). Importantly, immunostaining of L1 spinal cords demonstrated selective, nuclear accumulation of p53 and p73 in ∼20% of motor neurons from ChAT-Cre mice (Figure 5E-H). Furthermore, RT-qPCR analysis of p73 mRNA isoforms demonstrated strong induction of ΔNp73 mRNA but neither TAp73 nor total p73 mRNAs in the spinal cord of ChAT-Cre mice relative to controls without Cre at P13 (Figure 5I). Together, these results indicate that the deficiency of SMN in motor neurons induces p53 and ΔNp73 expression through cell autonomous mechanisms, which correlates with neurodegeneration.

**Figure 5.**
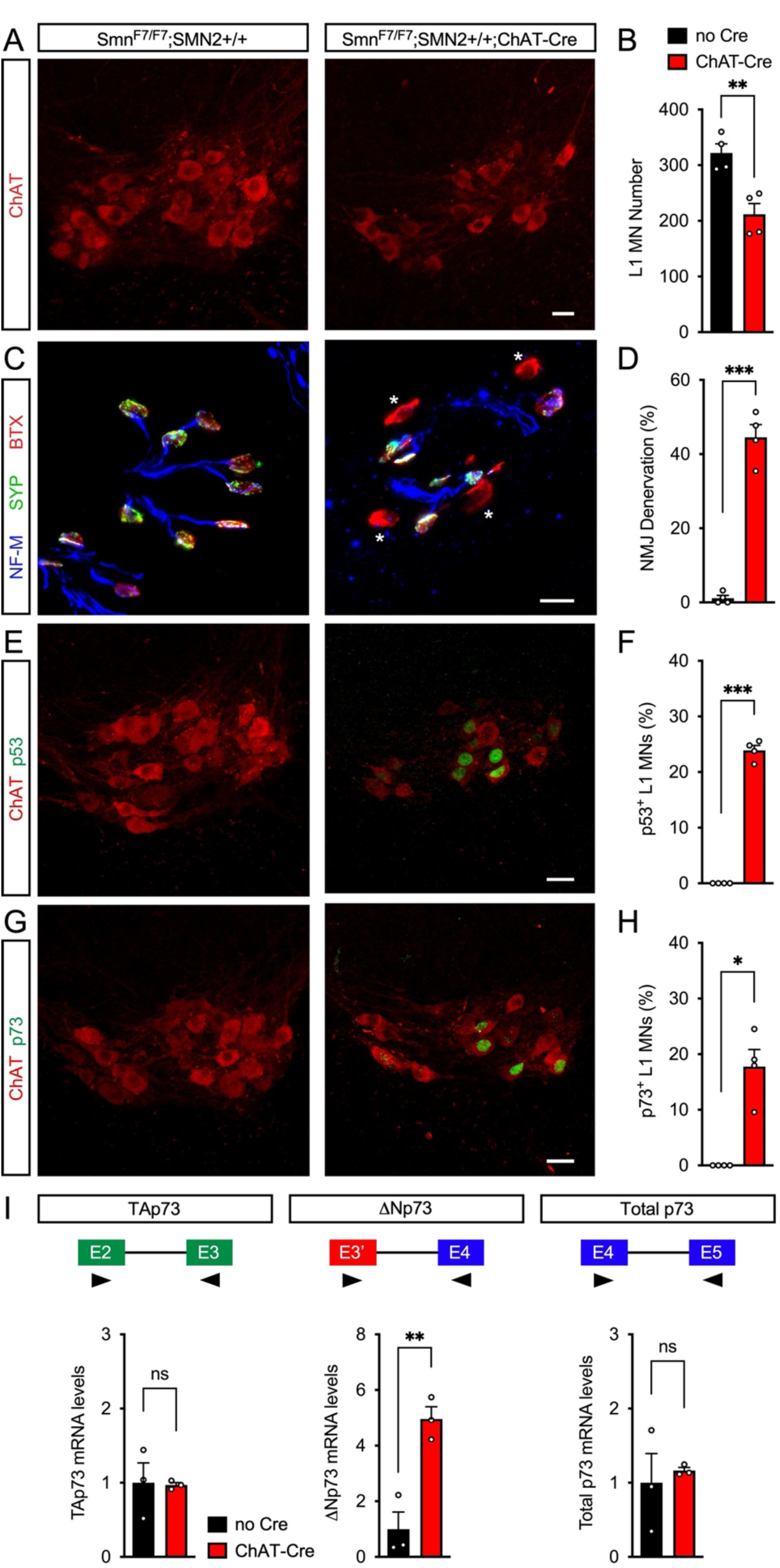
SMN deficiency drives cell autonomous p73 expression in motor neurons. (**A**) ChAT immunostaining of L1 spinal cords isolated at P13 from *Smn^F7/F7^; SMN2^+/+^* mice with or without the ChAT-Cre allele. Scale bar = 25 µm. (**B**) Total number of L1 motor neurons in the same experimental groups as in (A). The graph shows mean, SEM, and individual values from no Cre (n=4), and ChAT-Cre (n=4) mice. Two-tailed unpaired Student’s t-test. **P < 0.01. (**C**) NMJ staining with bungarotoxin (BTX), Synaptophysin (SYP), and Neurofilament-M (NF-M) of QL muscles isolated at P13 from the same experimental groups as in (A). Asterisks mark fully denervated NMJs. Scale bar = 25 µm. (**D**) Percentage of fully denervated NMJs in the QL muscle from experiments as in (C). The graph shows mean, SEM, and individual values from no Cre (n=4) and ChAT-Cre (n=4) mice. Welch’s test. ***P < 0.001. (**E**) ChAT and p53 immunostaining of L1 spinal cords isolated at P13 from the same experimental groups as in (A). Scale bar = 25 µm. (**F**) Percentage of p53^+^ L1 motor neurons from experiments as in (E). The graph shows mean, SEM, and individual values from no Cre (n=4) and ChAT-Cre (n=4) mice. Welch’s test. ***P < 0.001. (**G**) ChAT and p73 immunostaining of L1 spinal cords isolated at P13 from the same experimental groups as in (A). Scale bar = 25 µm. (**H**) Percentage of p73^+^ L1 motor neurons from experiments as in (G). The graph shows mean, SEM, and individual values from no Cre (n=4) and ChAT-Cre (n=4) mice. Welch’s test. *P < 0.05. (**I**) RT-qPCR analysis of TAp73, ΔNp73 and total p73 mRNAs in P13 spinal cords from the same experimental groups as in (A). The graph shows mean, SEM, and individual values normalized to no Cre samples as a control (n=3 mice). Two-tailed unpaired Student’s t-test for ΔNp73 mRNA, Welch’s test for TAp73 and total p73 mRNAs. **P < 0.01; ns, not significant. The schematics show the location of the PCR primers.

### SMN deficiency induces p73 expression through p53 and p38MAPK activation

To address the mechanisms of p73 induction in SMA motor neurons, we first tested its SMN dependency by analyzing the effects of adeno associated virus serotype 9 (AAV9)-mediated SMN restoration following intracerebroventricular (ICV) injection in neonatal SMA mice, which we previously showed robustly improves the disease phenotype^23^. Immunostaining of L1 spinal cords from untreated WT and SMA mice as well as SMA mice injected with AAV9-SMN at P0 demonstrated that SMN gene delivery almost completely abolished the expression of both p53 and p73 proteins in SMA motor neurons (Figure 6), confirming that their induction is dependent on SMN deficiency.

**Figure 6.**
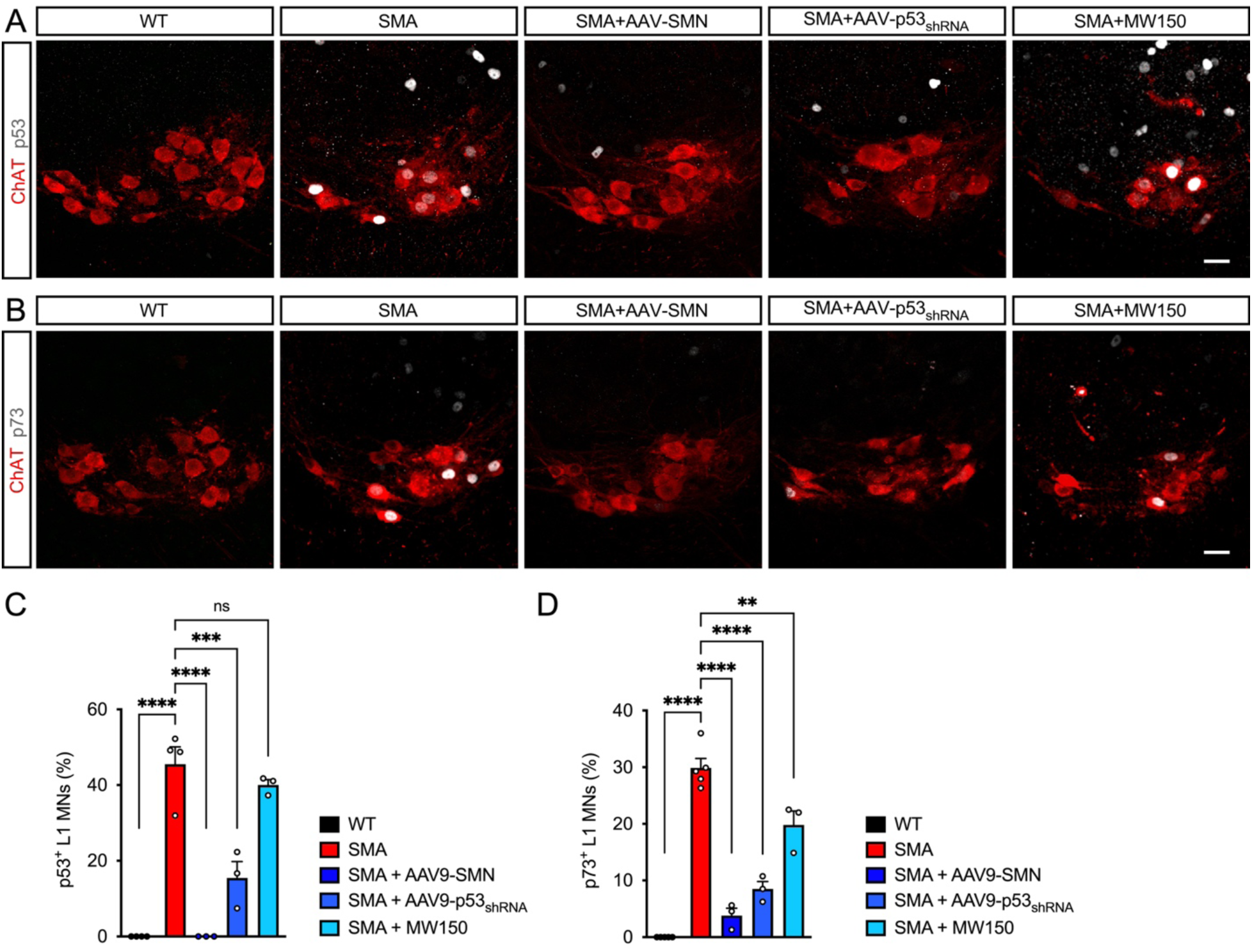
SMN deficiency induces p73 expression through p53 and p38MAPK activation in vulnerable SMA motor neurons. (**A**) ChAT and p53 immunostaining of L1 spinal cords isolated at P11 from uninjected WT and SMA mice, SMA mice injected with AAV9-SMN or AAV9-p53_shRNA_ at P0, and SMA mice treated daily with MW150 (5 mg/kg) from P0. Scale bar = 25 µm. (**B**) ChAT and p73 immunostaining of L1 spinal cords isolated at P11 from the same experimental groups as in (A). Scale bar = 25 µm. (**C**) Percentage of p53^+^ L1 motor neurons from experiments as in (A). The graph shows mean, SEM, and individual values from WT (n=4), SMA (n=4), SMA+AAV9-SMN (n=3), SMA+AAV9-p53_shRNA_ (n=3), and SMA+MW150 (n=3) mice. One-way ANOVA with Tukey’s *post hoc* test. ****P < 0.0001; ***P < 0.001; ns, not significant. (**D**) Percentage of p73^+^ L1 motor neurons from experiments as in (B). The graph shows mean, SEM, and individual values from WT (n=4), SMA (n=5), SMA+AAV9-SMN (n=3), SMA+AAV9-p53_shRNA_ (n=3), and SMA+MW150 (n=3) mice. One-way ANOVA with Tukey’s *post hoc* test. ****P < 0.0001; **P < 0.01.

To determine whether p53 activation drives p73 expression in SMA motor neurons, we used a previously validated AAV9 vector expressing a short hairpin RNA (AAV9-p53_shRNA_) targeting endogenous mouse p53 mRNA as well as GFP to monitor transduction efficiency (Supplementary Figure S4A)^13^. SMA mice were ICV injected at P0 with AAV9-p53_shRNA_ and L1 spinal segments isolated at P11 for analysis of p53 and p73 expression in motor neurons relative to uninjected WT and SMA mice as controls. Consistent with previous studies, analysis of GFP expression confirmed the high efficiency of transduction (∼70%) and the robust inhibition of p53 in SMA motor neurons by AAV9-p53_shRNA_ (Figure 6A and 6C, and Supplementary Figure S4B and S4C). Importantly, p53 knockdown also resulted in the strong reduction of the percentage of motor neurons expressing p73 in SMA mice (Figure 6B and 6D). These findings demonstrated that p53 activation is upstream of p73 and directly responsible for its expression in SMA motor neurons.

Given that the profile of p53 upregulation is much broader than that of p73 in the spinal cord of SMA mice, additional mechanisms must be at play that control p53 transcriptional activity to induce p73 expression selectively in vulnerable motor neurons. One possible candidate is phosphorylation of the amino terminal of p53 including serine 18 (p53^S18^), which we previously showed to be restricted to vulnerable SMA motor neurons and to contribute to their death^13^. Furthermore, we implicated p38α MAPK in the phosphorylation of p53^S18^ and showed that its inhibition is neuroprotective in SMA mice^13,24^. Unfortunately, we could not test by immunohistochemistry whether p73 accumulation occurs in the subset of SMA motor neurons expressing phosphorylated p53^S18^ because the commercial antibodies used in our previous studies are no longer available. Therefore, we analyzed the effects of pharmacological inhibition of p38α MAPK on p73 accumulation in motor neurons of SMA mice. As in our previous studies^13,24^, we treated SMA mice with daily IP injections of the highly selective inhibitor MW150 (5mg/kg) starting at birth and isolated spinal cords at P11 for immunostaining with p53 and p73. Interestingly, while MW150 treatment did not affect the expression of p53 as expected (Figure 6A and 6C), we found a significant reduction in the percentage of p73^+^ motor neurons in the L1 spinal cord from MW150 treated relative to untreated SMA mice (Figure 6B and 6D). These findings suggest that p38α MAPK activation partially contributes to the p53-dependent induction of p73 in SMA motor neurons.

### p73 drives motor neuron degeneration in SMA mice

To determine whether p73 induction contributes to the degeneration of SMA motor neurons, we employed an *in vivo* knockdown approach using an AAV9 vector (AAV9-p73_shRNA_) expressing a previously validated shRNA targeting all endogenous mouse p73 mRNA isoforms^47^. The AAV9-p73_shRNA_ vector also expresses GFP to monitor *in vivo* transduction (Supplementary Figure S4D). SMA mice were ICV injected with AAV9-p73_shRNA_ at P0 and L1 and L5 spinal cord segments were isolated at P11 for analysis of p53 and p73 expression in motor neurons relative to uninjected WT and SMA mice, which were used as controls. The analysis of GFP expression demonstrated high transduction efficiency of vulnerable L1 (∼70%) and L5 MMC (∼60%) SMA motor neurons (Supplementary Figure S4E-H). Importantly, the delivery of AAV9-p73_shRNA_ strongly inhibited the nuclear accumulation of p73, but had no effects on that of p53, in both L1 and L5 MMC motor neurons of SMA mice (Figure 7 and Supplementary Figure S5). These results demonstrated the effective and specific knockdown of p73 *in vivo* and confirmed our findings that p53 upregulation lies upstream of p73 induction in vulnerable SMA motor neurons.

**Figure 7.**
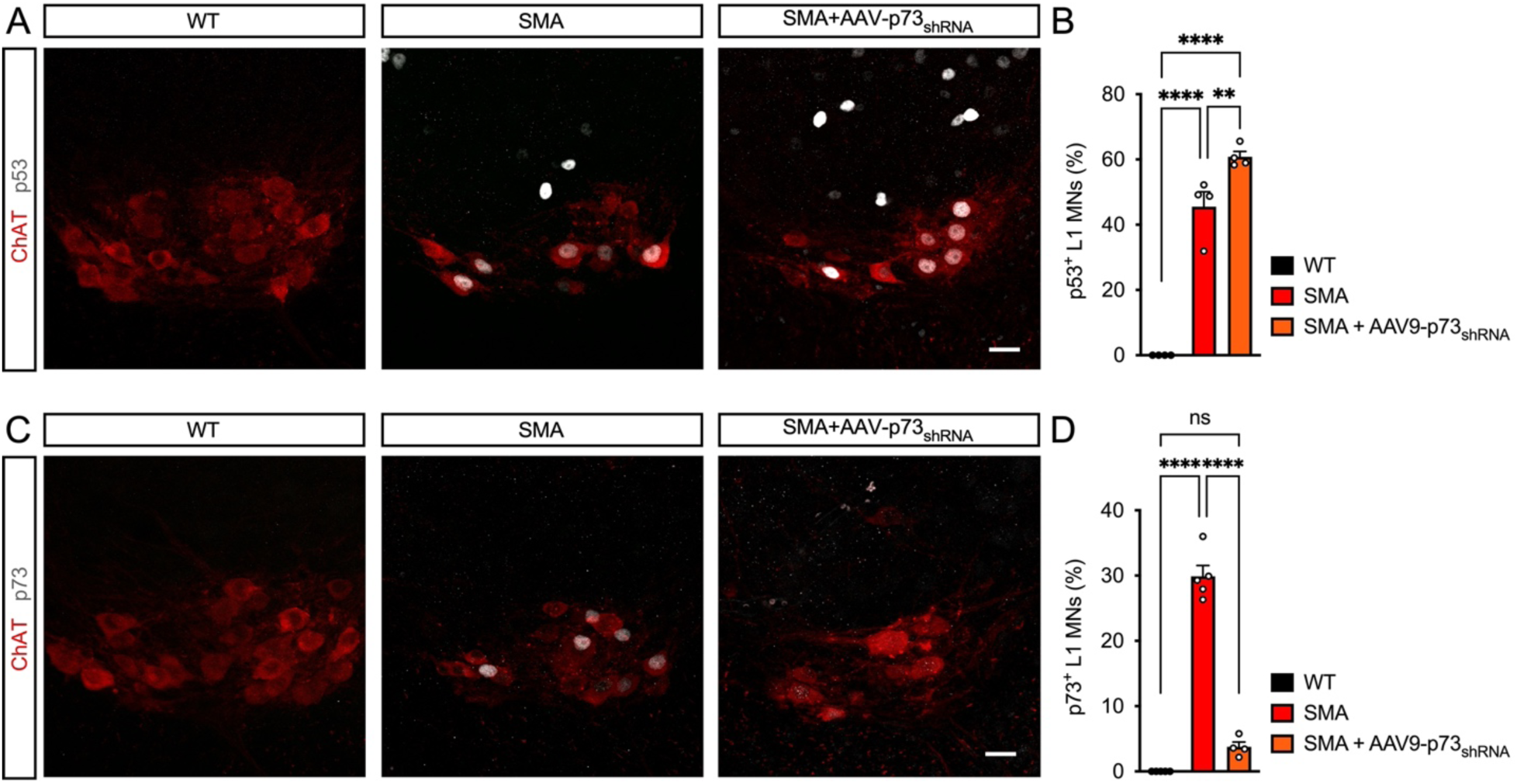
AAV9-mediated p73 knockdown does not affect p53 induction in motor neurons of SMA mice. (**A**) ChAT and p53 immunostaining of L1 spinal cords isolated at P11 from uninjected WT and SMA mice, and SMA mice injected with AAV9-p73_shRNA_ at P0. Scale bars = 25 µm. (**B**) Percentage of p53^+^ L1 motor neurons from experiments as in (A). The graph shows mean, SEM, and individual values from WT (n=4), SMA (n=4), and SMA+AAV9-p73_shRNA_ (n=4) mice. One-way ANOVA with Tukey’s *post hoc* test. ****P < 0.0001; **P < 0.01; ns, not significant. (**C**) ChAT and p73 immunostaining of L1 spinal cords isolated at P11 from uninjected WT and SMA mice, and SMA mice injected with AAV9-p73_shRNA_ at P0. Scale bars = 25 µm. (**D**) Percentage of p73^+^ L1 motor neurons from experiments as in (C). The graph shows mean, SEM, and individual values from WT (n=4), SMA (n=5), and SMA+AAV9-p73_shRNA_ (n=4) mice. One-way ANOVA with Tukey’s *post hoc* test. ****P < 0.0001.

We next investigated the effects of blocking p73 induction on motor neuron degeneration in SMA mice. To ensure that any potential effects are independent of SMN upregulation, we analyzed SMN protein levels in the spinal cord of SMA mice injected with AAV9-p73_shRNA_ relative to uninjected WT and SMA mice as controls. Western blot analysis showed that the low levels of SMN found in the spinal cord of untreated SMA mice at P11 were not affected by the injection of AAV9-p73_shRNA_ and the expression of GFP confirmed robust tissue transduction (Figure 8A and 8B). Importantly, analysis of vulnerable motor neuron pools in the same experimental groups revealed significantly increased numbers of L1 and L5 MMC motor neurons in SMA mice treated with AAV9-p73_shRNA_ relative to uninjected SMA mice (Figure 8C-E). Consistent with previous studies^12,13^, the neuroprotective effects were less pronounced in L1 motor neurons likely due to the earlier onset and more rapid progression of the degenerative process in this vulnerable motor neuron pool.

**Figure 8.**
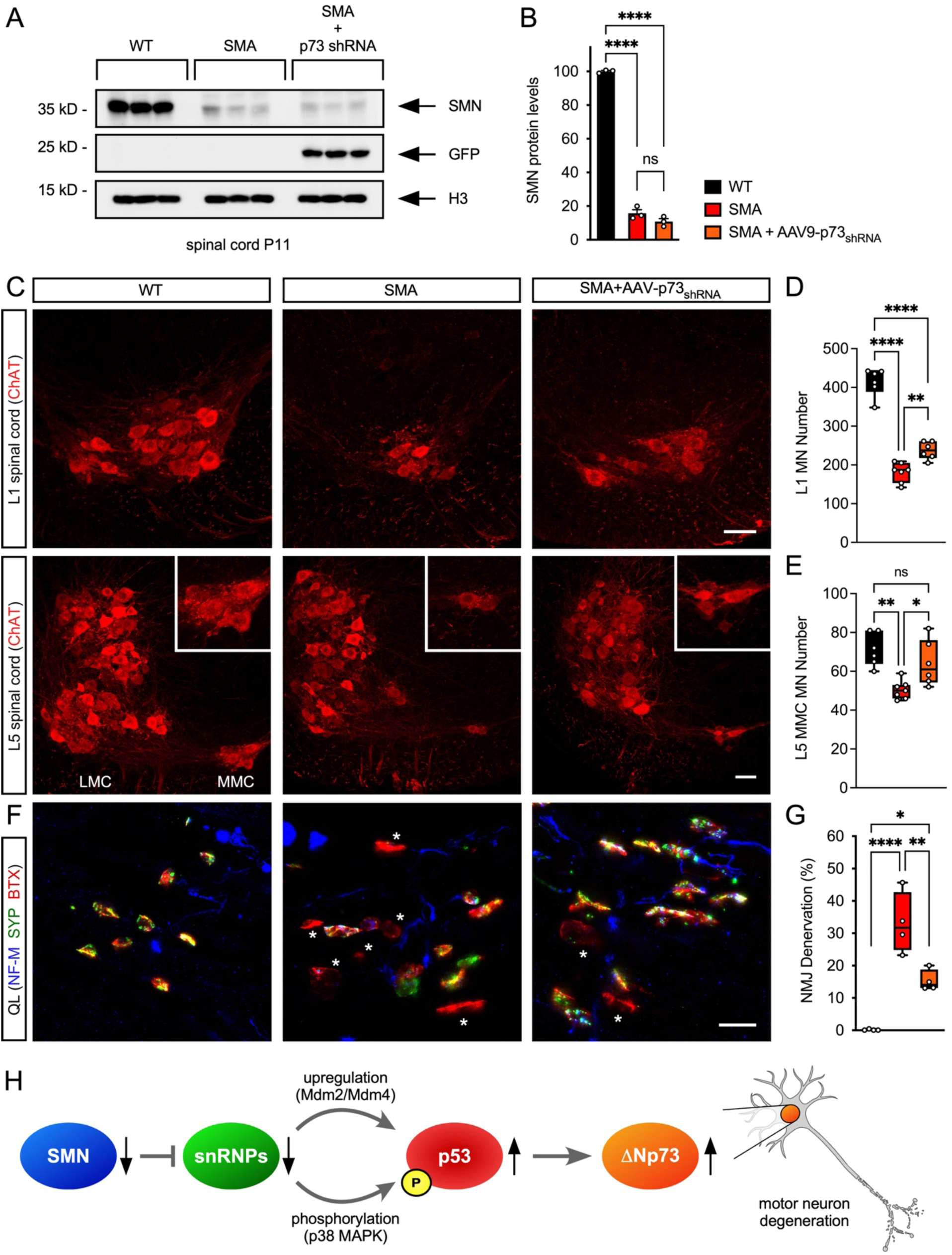
p73 drives motor neuron degeneration in SMA mice. (**A**) Western blot analysis of SMN and GFP expression in P11 spinal cords from uninjected WT and SMA mice, and SMA mice injected with AAV9-p73_shRNA_ at P0. Histone H3 was used as loading control. (**B**) Quantification of SMN levels from the experiment in (A). Normalized mean, SEM, and individual values from 3 independent biological replicates are shown. One-way ANOVA followed by Tukey’s *post hoc* test. ****P < 0.0001; ns, not significant. (**C**) ChAT immunostaining of L1 and L5 spinal cords isolated at P11 from uninjected WT and SMA mice, and SMA mice injected with AAV9-p73_shRNA_ at P0. L5 LMC and MMC motor neuron pools are indicated, and magnified views of L5 MMC motor neurons are shown in the insets. Scale bars = 50 µm. (**D**) Total number of L1 motor neurons from experiments as in (C). The box and whiskers graph shows the median, interquartile range, minimum, maximum and individual values from WT (n=6), SMA (n=7), and SMA+AAV9-p73_shRNA_ (n=6) mice. One-way ANOVA with Tukey’s *post hoc* test. ****P < 0.0001; **P < 0.01. (**E**) Total number of L5 MMC motor neurons from experiments as in (C). The box and whiskers graph shows the median, interquartile range, minimum, maximum and individual values from WT (n=6), SMA (n=7), and SMA+AAV9-p73_shRNA_ (n=6) mice. One-way ANOVA with Tukey’s *post hoc* test. **P < 0.01; *P < 0.05; ns, not significant. (**F**) NMJ staining with bungarotoxin (BTX), Synaptophysin (SYP), and Neurofilament-M (NF-M) of QL muscles isolated at P11 from uninjected WT and SMA mice, and SMA mice injected with AAV9-p73_shRNA_ at P0. Asterisks mark fully denervated NMJs. Scale bar = 25 µm. (**G**) Percentage of fully denervated NMJs in the QL muscle from experiments as in (F). The box and whiskers graph shows the median, interquartile range, minimum, maximum and individual values from WT (n=4), SMA (n=4), and SMA+AAV9-p73_shRNA_ (n=4) mice. One-way ANOVA followed by Tukey’s *post hoc* test. ****P < 0.0001; **P < 0.01; *P < 0.05. (**H**) Schematic model of the events induced by SMN deficiency that lead to ΔNp73 expression and selective neurodegeneration of vulnerable motor neurons in SMA. SMN deficiency disrupts snRNP assembly, resulting in defective splicing of specific genes that cause p53 upregulation (Mdm2/Mdm4) and its amino-terminal serine 18 phosphorylation by p38 MAPK (Stasimon). These converging events, possibly in combination with yet-to-be-identified modifications, promote isoform-specific expression of ΔNp73, leading to the selective death of vulnerable SMA motor neurons.

Lastly, we analyzed the effects of p73 inhibition on neuronal innervation of the QL muscle by immunostaining of presynaptic (neurofilament and synaptophysin) and postsynaptic (bungarotoxin labeling of acetylcholine receptor clusters) compartments. We found a robust, yet incomplete reduction in the percentage of denervated NMJs in SMA mice treated with AAV9-p73_shRNA_ (Figure 8F and 8G), which aligns well with the degree of rescue of L1 motor neurons.

Collectively, these results link p53-mediated, selective expression of ΔNp73 in vulnerable SMA motor neurons to the execution of the intrinsic neurodegenerative process induced by SMN deficiency.

## Discussion

Selective motor neuron degeneration is a defining feature of SMA, yet the mechanisms linking ubiquitous SMN deficiency to highly restricted neuronal loss have remained incompletely understood. Previous studies established that SMN deficiency activates p53 signaling in SMA motor neurons^13^, but how this pathway is selectively converted into a degenerative program in vulnerable but not resistant neurons remained unresolved. Here, we identify ΔNp73 as a selective downstream effector of p53-dependent degeneration in SMA. We show that ΔNp73 is induced specifically in vulnerable motor neuron populations in both SMA mice and type 1 SMA patients, depends on p53 activation, and functionally contributes to motor neuron loss and neuromuscular denervation. Thus, we uncover a p53–ΔNp73 signaling axis that links SMN deficiency to selective neurodegeneration in SMA.

Our findings substantially refine the current molecular framework of selective motor neuron degeneration in SMA (Figure 8H). Prior work established that SMN-dependent disruption of snRNP assembly initiates convergent RNA-processing defects leading to stabilization of p53 and its phosphorylation by p38 MAPK in vulnerable SMA motor neurons^13,22–24^. While these studies established p53 as a central mediator of degeneration, they did not explain how p53 activation is translated into selective neuronal death. The identification of ΔNp73 as a downstream component of this pathway provides a mechanistic bridge between p53 activation and the execution phase of degeneration. Within this context, selective transcriptional reprogramming downstream of p53 leads to induction of ΔNp73, which acts as a critical downstream contributor to neurodegeneration. Furthermore, the observation that selective depletion of SMN in motor neurons is sufficient to induce p53 and ΔNp73 expression demonstrates that this degenerative pathway arises through neuron intrinsic mechanisms, providing a molecular signature of the cell autonomous signaling cascade coupling SMN deficiency to selective motor neuron degeneration^44,45^. Importantly, our findings suggest that selective neuronal vulnerability in SMA is determined by context-dependent reprogramming of p53 family signaling downstream of SMN deficiency. Accordingly, p53 accumulation alone is insufficient to determine neuronal death and extends beyond vulnerable SMA motor neurons at later stages of disease^13,22^. In contrast, ΔNp73 induction remains highly restricted, indicating that additional regulatory mechanisms shape the transcriptional output of p53 selectively within susceptible neuronal populations. One plausible mechanism involves phosphorylation of p53 mediated by p38 MAPK, which we previously showed to occur selectively in vulnerable SMA motor neurons and contribute to degeneration^13,23,24^. Consistent with this, we find that pharmacological inhibition of p38 MAPK reduces ΔNp73 induction without altering overall p53 accumulation, supporting the idea that p38 MAPK-dependent signaling influences p53 target selection, including ΔNp73 expression in vulnerable neurons. However, the partial effect of p38 MAPK inhibition also suggests that additional regulatory layers cooperate to direct the selective transcriptional program associated with degeneration.

This study reveals that ΔNp73 upregulation promotes degeneration in SMA motor neurons, a role that stands in stark contrasts to its established functions in cancer and neurodevelopment^34,38,39^. ΔNp73 is known to function as a dominant negative inhibitor of p53- and TAp73-mediated transcription^33^ as well as a factor that promotes neuronal survival in development^48,49^. Induction of ΔNp73 would therefore be predicted to oppose p53 dependent neuronal death in SMA. Instead, our data indicate that ΔNp73 induction drives degeneration, revealing a striking functional switch in diseased motor neurons. These findings suggest that the output of p53-ΔNp73 signaling is highly context-dependent and may differ fundamentally across developmental, oncogenic, and neurodegenerative settings. Leveraging both p53-dependent and p53-independent functions of ΔNp73 in gene regulation^33,50–52^, this degenerative shift may reflect altered promoter selection, interactions with additional transcription factors, and post translational modification of p53-family proteins in vulnerable SMA motor neurons. In this context, p38 MAPK may contribute not only through phosphorylation of p53 but also of p73^53^. Furthermore, we found that PADI4 is strongly upregulated (Supplementary Table S1), and protein citrullination emerged among the most dysregulated pathways in SMA motor neurons (Figure 1F). Recent evidence demonstrating that PADI4-mediated citrullination can redirect p53 transcriptional activity^54^ raises the possibility that similar mechanisms shape p53-ΔNp73 signaling toward a degenerative program selectively in vulnerable SMA motor neurons.

The identification of ΔNp73 as a downstream effector of neurodegeneration further supports a model in which SMA motor neuron death proceeds through non-canonical p53-dependent mechanisms, which is consistent with the lack of caspase activation and DNA fragmentation^13^. Rather than activating a classical apoptotic program, p53-ΔNp73 signaling may engage alternative cell death pathways that drive neuronal degeneration. Defining these downstream programs will be important for understanding the molecular execution of degeneration and may reveal broader principles governing neuronal death pathways. Interestingly, our transcriptional profiling identified additional genes that are dysregulated in an SMN- and p53-dependent manner and may contribute to downstream degenerative processes. In particular, reduced expression of Atp11c and Atp2b4 (Figure 1 and Supplementary Figure S1) suggests that altered phospholipid homeostasis and membrane asymmetry may participate in neuronal loss. Atp11c regulates phosphatidylserine membrane asymmetry through its flippase activity, whereas Atp2b4 controls calcium extrusion which is linked to phosphatidylserine exposure through regulation of plasma membrane scramblases^55,56^. Disruption of these pathways along the p53-ΔNp73 axis could expose “eat-me” signals that promote phagocytic removal of SMA neurons.

Beyond their mechanistic significance, our findings also have potential therapeutic implications. Current SMA therapies restore SMN expression but do not fully prevent neurodegeneration or reverse established motor deficits, particularly after symptom onset^30–32^. These limitations highlight the need for complementary neuroprotective strategies targeting downstream pathogenic pathways. Although direct inhibition of p53 is unlikely to be clinically viable due to its essential tumor suppressor functions, targeting downstream effectors may offer greater selectivity. In this context, selective inhibition of ΔNp73 through isoform-specific antisense approaches could provide neuroprotection while preserving tumor suppressive functions mediated by TAp73^57^. As shown with p38 MAPK inhibition^24^, combinatorial interventions targeting neuron intrinsic death pathways may extend therapeutic benefit beyond that achievable through SMN augmentation alone.

While this study focuses on spinal motor neurons, degeneration or dysfunction of additional neuronal populations contributes to SMA pathology. Recent studies have reported loss of locus coeruleus neurons^58,59^ and cerebellar Purkinje cells^60^ in SMA mouse models, and early evidence suggests that p53 signaling also contributes to Purkinje cell degeneration^60^. It will therefore be important to determine whether the p53-ΔNp73 signaling axis identified here represents a broader mechanism of SMN-dependent neurodegeneration. More generally, our findings may have implications beyond SMA. p53 signaling has been implicated in a range of neurodegenerative disorders^15^, including amyotrophic lateral sclerosis^16–19^, Huntington’s disease^61,62^, Parkinson’s disease^63–65^, and Alzheimer’s disease^66–68^, as well as neuronal damage following ischemic stroke and traumatic brain injury^69,70^. The mechanisms described here may therefore reflect broader principles by which p53 transcriptional pathways are selectively reprogrammed to drive neuronal vulnerability in disease.

In summary, we identify ΔNp73 as a selective downstream effector of p53-dependent motor neuron degeneration in SMA and demonstrate that its inhibition mitigates neuronal loss and neuromuscular denervation *in vivo*. These findings reveal a previously unrecognized, context dependent reprogramming of p53 family signaling in neurodegeneration and provide new mechanistic insight into how broadly expressed cellular stress pathways produce highly selective neuronal vulnerability. More broadly, our study suggests that neuronal susceptibility in neurodegenerative disease may arise not simply from general activation of stress pathways themselves, but from cell type-specific interpretation of those signals through selective transcriptional programs.

## Methods

### DNA constructs and AAV9 viral vectors

AAV9-p53_shRNA_ and AAV9-SMN vectors were previously described^13,23^. A p73 shRNA cassette driven by the H1 promoter was synthesized by Genewiz based on a previously published sequence (5′-GGGACTTCAATGAAGGACA-3′) targeting mouse *Trp73*^47^. The cassette was cloned into an AAV9 vector harboring a GusB promoter driving GFP expression in the opposite orientation using standard molecular cloning techniques. The corresponding AAV9 viral vector was custom produced by VectorBuilder, and viral titer and purity were confirmed in-house by qPCR using primers targeting the BGH polyadenylation sequence (Supplementary Table S3) and by silver staining, respectively. Expression plasmids encoding HA-tagged mouse ΔNp73 (NM_011642.4) or TAp73 (NM_001126330.1) under the control of the CBA promoter were generated by VectorBuilder. All constructs were verified by Sanger DNA sequencing.

### Cell lines and treatments

HeLa CCL-2 cells and HEK293T cells were obtained from the American Type Culture Collection (ATCC) without further authentication. All cell lines were cultured in Dulbecco’s Modified Eagle’s Medium (DMEM) with high glucose (Gibco) containing 10% fetal bovine serum (HyClone), 2 mM L-glutamine (Gibco), and 0.1mg/ml gentamicin (Gibco). Cells were regularly tested for mycoplasma using the Universal Mycoplasma Detection Kit (ATCC). HEK293T and HeLa CCL-2 cells were transfected with HA-tagged ΔNp73 or HA-TAp73 expression plasmids using PEI MAX (Polysciences) at a DNA:PEI ratio of 1:2 according to the manufacturer’s instructions.

### Human tissue

Human spinal cords were collected through autopsy procedures performed at the Johns Hopkins University, Baltimore, MD, USA following parental- or patient-informed consent in strict observance of the legal and institutional ethical regulations. None of the SMA patients received disease-modifying therapies and further information is provided in Supplementary Table S2.

### Mouse lines

All mouse work was performed in accordance with the National Institutes of Health Guidelines on the Care and Use of Animals, complied with all ethical regulations and was approved by the Institutional Animal Care and Use Committee (IACUC) of Columbia University (AABR9600). Mice were housed in a 12h/12h light/dark cycle with access to food and water *ad libitum*. All original breeding pairs were obtained from The Jackson Laboratory. FVB.Cg-*Grm7^Tg(SMN2)89Ahmb^ Smn1^tm1Msd^* Tg(SMN2*delta7)4299Ahmb/J (JAX #005025) mice were interbred to obtain SMA mutant mice^71^. Mutant Smn^F7/F7^; SMN2^+/+^; ChAT-Cre mice were generated by crossing ChAT-Cre mice (B6;129S6-Chat^tm2(cre)Lowl^/J; JAX #006410) with mice carrying the floxed Smn^F7^ Cre-inducible knockout allele and the human SMN2 transgene (Cg-Grm7^Tg(SMN2)89Ahmb^ Smn1^tm1Jme^/J; JAX #033847) followed by selection of Cre-positive offspring and backcrossing to obtain the experimental genotype. Genotyping was performed from tail DNA as previously described for the SMNΔ7^72^ and *Smn^F^*^743^ mouse lines. Equal proportions of mice of both sexes were used and aggregated data are presented because sex-specific differences were not found. A list of the primers used is shown in Supplementary Table S3.

### Animal procedures

For drug delivery, SMA mice were treated daily via IP injection starting at P0 with SMN-C3 (3 mg/kg) or Pifithrin-α (2.2 mg/kg) in DMSO, or MW150 (5 mg/kg) in saline, as previously described^13,24^. For AAV9-mediated gene delivery, ICV injections were carried out at P0 in SMNΔ7 mice anesthetized by isoflurane inhalation by a single injection in the right lateral ventricle of the brain of ∼1×10^11^ genome copies of AAV9 vectors in 5 µl of a phosphate buffered saline (PBS) solution containing 5% sorbitol, 0.001% Pluronic F-68, and Fast Green dye (Sigma). For muscle injections, anesthesia was induced and maintained with isoflurane. Mice were placed in the supine position, and a small unilateral abdominal incision was made under a stereomicroscope (Zeiss). The abdominal wall was gently retracted, and the iliopsoas muscle was identified beneath the intact peritoneum. A physiological saline solution containing 1% Alexa Fluor 488–conjugated cholera toxin subunit B (CTB-488, Thermo Fisher Scientific) was injected into the IL and QL muscles of experimental mice at P2 using a 5 µl Hamilton syringe. Skin was closed with microsurgical nylon sutures (Fine Science Tools).

### Laser capture microdissection

Mice were euthanized at P6, and spinal cords were collected under continuous oxygenation in cold artificial cerebrospinal fluid (aCSF). Successful tracer delivery was confirmed by visualization of fluorescent signal in the injected muscle and corresponding motor neurons. Lumbar spinal cord segments L1–L3 were dissected, immediately embedded in OCT compound (Fisher Healthcare Tissue-Plus), snap-frozen on dry ice, and stored at −80°C. Frozen tissue was cryosectioned longitudinally (14 µm), mounted onto UV-treated PEN membrane slides (Applied Biosystems), and stored at −80°C until LCM was performed. Immediately prior to LCM, sections were fixed in 70% ethanol for 10 s and dehydrated sequentially in 95% ethanol (10 s, twice) and 100% ethanol (30–60 s, twice), followed by air-drying. LCM was performed using a Leica DM6000B microscope under 20X magnification to isolate CTB-488–positive motor neurons from L1–L3 spinal cord sections of control and experimental mice. Approximately 600 motor neurons were collected from L1–L3 spinal segments from one or more mice for each biological replicate, and captured cells were collected into tube caps containing 50 µl lysis buffer to preserve RNA integrity. RNA was extracted from LCM-isolated neurons using the Absolutely RNA Nanoprep Kit (Agilent Technologies), including on-column DNase treatment, according to the manufacturer’s instructions. RNA quality was assessed using an Agilent 2100 Bioanalyzer with the Eukaryote Total RNA Pico assay.

### RNA sequencing and bioinformatics

RNA sequencing libraries were prepared and sequenced at the Columbia Sulzberger Genome Center. Total RNA isolated from LCM motor neurons (<10ng per sample) was processed using a low input polyadenylated RNA amplification and library preparation workflow. Libraries were normalized by the core facility and sequenced on an Illumina NovaSeq platform (paired-end 100 bp; ∼40M reads per sample). A total of 15 libraries were sequenced (WT, SMA, SMA+PFTα, SMA+MW150, SMA+SMN-C3; n=3 biological replicates per group). Post-sequencing processing and primary data analysis were performed by the Columbia Sulzberger Genome Center. Reads were pseudoaligned to the mouse transcriptome (Ensembl GRCm38.p6) using kallisto (v0.44.0) to generate transcript- and gene-level abundance estimates. Differential gene expression analysis between experimental groups was performed using DESeq2 with standard parameters. Isoform transcript quantification was done using RSEM (v1.3.3) pipeline. Mouse reference transcriptome was built from GRCm38.p6 genome and GENCODE vM23 annotation. Bowtie2 (v2.5.1) was used for alignment transcript quantification. Indexing of sorted BAM files was done using Samtools (v1.18) and CPM normalized Bigwig tracks were generated using deepTools.

### RNAscope combined with immunofluorescence

Spinal cords of P6 WT or SMA mice were fixed by transcardial perfusion with 4% paraformaldehyde (PFA) in PBS, dissected, and post-fixed in 4% PFA in PBS for 24 h at 4°C. L1 spinal cord segments were identified by nerve roots, dissected, cryoprotected in 10%, 20%, and 30% sucrose in phosphate buffer at 4°C, embedded in OCT compound, and frozen on dry ice. Cryosections (20μm) were cut onto Superfrost Plus slides, washed in PBS for 5 min to remove OCT, and post-fixed in 4% PFA in PBS for 60 min. Sections were dehydrated in 50%, 70%, and 100% ethanol (twice) for 5 min each at RT. RNAscope was performed using the RNAscope Multiplex Fluorescent Reagent Kit v2 with TSA Vivid Dye (Bio-Techne/ACD) following the manufacturer’s instructions for manual workflow. Briefly, sections were circled with a hydrophobic pen, treated with RNAscope Hydrogen Peroxide for 10 min at RT, rinsed in distilled water, and incubated with RNAscope Protease Plus for 20 min at 40°C. RNAscope probes were hybridized for 2 h at 40°C (Supplementary Table S4). Sections were washed and stored overnight in 5x SSC at RT. The following day, sections were sequentially incubated with AMP1, AMP2 (30 min), and AMP3 (15 min) at 40°C, with washes between steps. Signal was developed using HRP (15 min), TSA fluorophore (30 min) and HRP blocking (15 min) at 40°C with washes between steps. For immunofluorescence, sections were washed in TBST, blocked with 10% normal donkey serum in TBS containing 1% BSA for 30 min at RT, and incubated with primary antibodies diluted in TBS-1% BSA overnight at RT. After washing, sections were incubated with species-appropriate secondary antibodies for 1 h at RT, washed, counterstained with DAPI for 30 s, and mounted with ProLong Gold Antifade Mountant (Invitrogen).

### RNA and Protein analysis

Mice were euthanized and spinal cords were collected in a dissection chamber under continuous oxygenation in cold aCSF, snap-frozen, and stored at −80°C until use. For RT–qPCR analysis, total RNA from whole spinal cord tissue was extracted with TRIzol reagent (Invitrogen) and treated with DNase I (Ambion). RNA was reverse transcribed using the RevertAid first-strand cDNA kit (Thermo Scientific). Quantitative PCR (qPCR) was carried out using Power SYBR green PCR master mix (Applied Biosystems) in QuantStudio 3 (Applied Biosystems). A list of the primers used is shown in Supplementary Table S3. For Western blot analysis, total protein extracts were generated by homogenization of spinal cords or cell pellets in SDS sample buffer (2% SDS, 10% glycerol, 5% Δ-mercaptoethanol, 60 mM Tris-HCl pH 6.8, and bromophenol blue), followed by brief sonication and boiling. Proteins were quantified using the *RC DC*^TM^ Protein Assay (Bio-Rad) and analyzed by SDS/PAGE on 12% polyacrylamide gels followed by Western blotting as previously described^73^. A list of the antibodies used is shown in Supplementary Table S5.

### Immunocytochemistry

HeLa CCL-2 cells were grown on coverslips and fixed in 4% paraformaldehyde. Following fixation, cells were washed with PBS, permeabilized with 0.1% Triton X-100 in PBS for 10 min, washed with PBS, and blocked in 1% BSA and 0.1% Tween-20 in PBS. Cells were incubated with primary antibodies in blocking buffer for 1 h at RT and with secondary antibodies in PBS for 1 h in the dark, with PBS washes between steps, followed by counterstaining with DAPI. Images were acquired using a confocal microscope (Leica).

### Immunohistochemistry of mouse tissue

For morphological studies by immunohistochemistry, mice were deeply anesthetized using Avertin, the depth of anesthesia was checked by the toe pinch reflex, and transcardial perfusion was then performed with a saline solution followed by 4% paraformaldehyde (PFA). The spinal cord and skeletal muscles were dissected and post-fixed by immersion in 4% PFA overnight at 4°C. For immunohistochemistry, the spinal cords were briefly washed with PBS and the L1 and L5 lumbar segments were identified by the ventral roots, dissected, and subsequently embedded in warm 5% agar. Transverse sections (75 μm) of the entire spinal segment were obtained with a VT1000 S vibratome (Leica). Free-floating sections were then incubated overnight at RT with different combinations of primary antibodies diluted in PBS-T. The following day, six washing steps with PBS-T of 10 min each were done prior to incubation with secondary antibodies for 3 h in PBS. Another six washing steps with PBS were performed before sections were mounted in 30% glycerol/PBS. For NMJ analysis, skeletal muscles were cryoprotected through sequential immersion in 10% and 20% sucrose/0.1M phosphate buffer (PB) for 1 h at 4°C followed by overnight immersion in 30% sucrose/0.1M PB at 4°C. The following day, muscles were embedded in OCT, frozen on dry ice, and stored at -80°C until processing. Longitudinal cryosections (30μm) were collected onto Superfrost Plus glass slides (Fisher Scientific) using a CM3050S cryostat (Leica). Sections were washed once with PBS for 5 min to remove OCT, blocked for 1 h with 5% donkey serum in TBS containing 0.2% Triton-X at RT and incubated with primary antibodies in blocking buffer overnight at 4°C. Following incubation, sections were washed three times for 10 min in TBS-T and then incubated with tetramethylrhodamine-conjugated α-bungarotoxin (Invitrogen) and the appropriate secondary antibodies for 1 h at RT, followed by 3 washing steps. Slides were mounted using Fluoromount-G Mounting Medium (SouthernBiotech). A list of the antibodies used is shown in Supplementary Table S5.

### Immunohistochemistry of human spinal cord

For immunohistochemistry on human spinal cord tissue, samples were first cryoprotected in 15% sucrose solution for ≥2 h until they sank, then moved to 30% sucrose and kept at 4°C overnight. The following day, tissue was mounted in Sakura Tissue-Tek OCT compound and rapidly frozen using 2-methylbutane chilled by liquid nitrogen. Sections were sliced at 20 µm thickness on a Leica CM3050 S cryostat at −20°C and stored at −80°C. Slides underwent antigen retrieval in L.A.B. solution (Polyscience) for 20 min at room temperature, followed by three PBS washes and blocking with 5% normal donkey serum in 0.3% PBS-T for 90 min. Primary antibodies were applied overnight at 4°C, then after three 10 min PBS rinses, secondary antibodies (diluted 1:1000 in PBS-T) were added for 3 h. Finally, following three 10 min PBS washes slides were coverslipped with a mounting medium containing glycerol:PBS (3:7). A list of the antibodies used is shown in Supplementary Table S5.

### Image analysis

Images were acquired using Leica SP8 or Stellaris confocal microscopes, or a Leica THUNDER DMi8 microscope under identical acquisition settings across experimental groups and analyzed offline using Leica LAS X software or ImageJ2 (Fiji, v2.9.0/1.53t). For RNAscope, signal was quantified using ImageJ2/Fiji according to the manufacturer’s recommended quantification guidelines. Individual ChAT⁺ motor neuron somata were manually outlined as regions of interest (ROIs). RNAscope puncta within each ROI were identified using automated intensity-based thresholding. Dot density was calculated as the number of RNAscope puncta normalized to the soma area (dots/µm²) for each motor neuron. Individual motor neurons were treated as technical replicates for quantification. For motor neuron quantification, confocal images were acquired from all 75 μm sections of L1 and L5 spinal segments using a 20X objective with 3μm z-steps. Only ChAT-positive motor neurons with a clearly identifiable nucleus were counted to avoid double counting from adjoining sections. p53 and p73 signals were evaluated on a binary basis (presence or absence of nuclear signal), to determine the percentage of positive cells. GFP was used to identify transduced cells. For NMJ analysis, images were acquired using a THUNDER DMi8 (Leica) microscope from 30 μm muscle sections using a 20X objective with software-optimized z-steps. Maximum intensity projections of stacks were analyzed and at least 200 randomly selected NMJs per muscle were quantified. NMJs lacking any coverage of the α-bungarotoxin-labeled postsynaptic endplate by the presynaptic markers Synaptophysin and Neurofilament-M were scored as denervated and those with less than 50% coverage were scored as partially innervated.

### Statistical analysis

Experimental animals were randomly assigned to treatment groups. The sample size for each experiment is detailed in the Figure legends and was determined based on previous publications. Results are expressed as mean and standard error of the mean (SEM) from at least three independent biological replicates unless otherwise indicated. The exact value and meaning of n for each dataset can be found in the Figure legends. The Shapiro-Wilk test was used to assess data normality. For normally distributed data, two groups were compared using a two-tailed unpaired Student’s t-test for equal variances or Welch’s test for unequal variances. The Mann-Whitney test was applied when normality assumptions were violated. For comparing three or more groups, a one-way ANOVA followed by Tukey’s *post-hoc* test was used for one categorical independent variable, while two-way ANOVA followed by Sidak’s multiple comparison test was used for two categorical independent variables. GraphPad Prism 11 for macOS Version 11.0.2 was used for all statistical analyses, and P values are indicated as follows: *P<0.05; **P<0.01; ***P<0.001; **** P<0.0001.

## Acknowledgements

We thank Daniel Martin Watterson for the kind gift of MW150. We are grateful to George Mentis for helpful discussions and critical reading of the manuscript. This work was supported by grants NS102451 (L.P.), NS114218 (L.P.) and NS116400 (L.P.) from NINDS, DFG Research Training Group GRK 3102 (C.M.S.), and DFG 19693-1 (C.M.S.). This research was funded in part through the NIH/NCI Cancer Center Support Grant P30CA013696 and used the Genomics and High Throughput Screening Shared Resource.

## Data availability

Sequencing data have been deposited in the Gene Expression Omnibus database under accession number GSE320428.

## Authors contributions

L.P. designed and supervised the study. M.J.C. performed and analyzed most experiments. X.H., M.S. and J.Q.G. contributed to experiments and data analysis. A.Y. and V.M. contributed to RNA-seq data analysis. C.J.S. provided human spinal cords. L.S. and C.M.S. performed immunostaining of human spinal cords. M.J.C and L.P. wrote the manuscript with input from all authors.

## Declaration of competing interest

The authors declare that they have no conflict of interest.

---

**Supplementary Figure S1.**
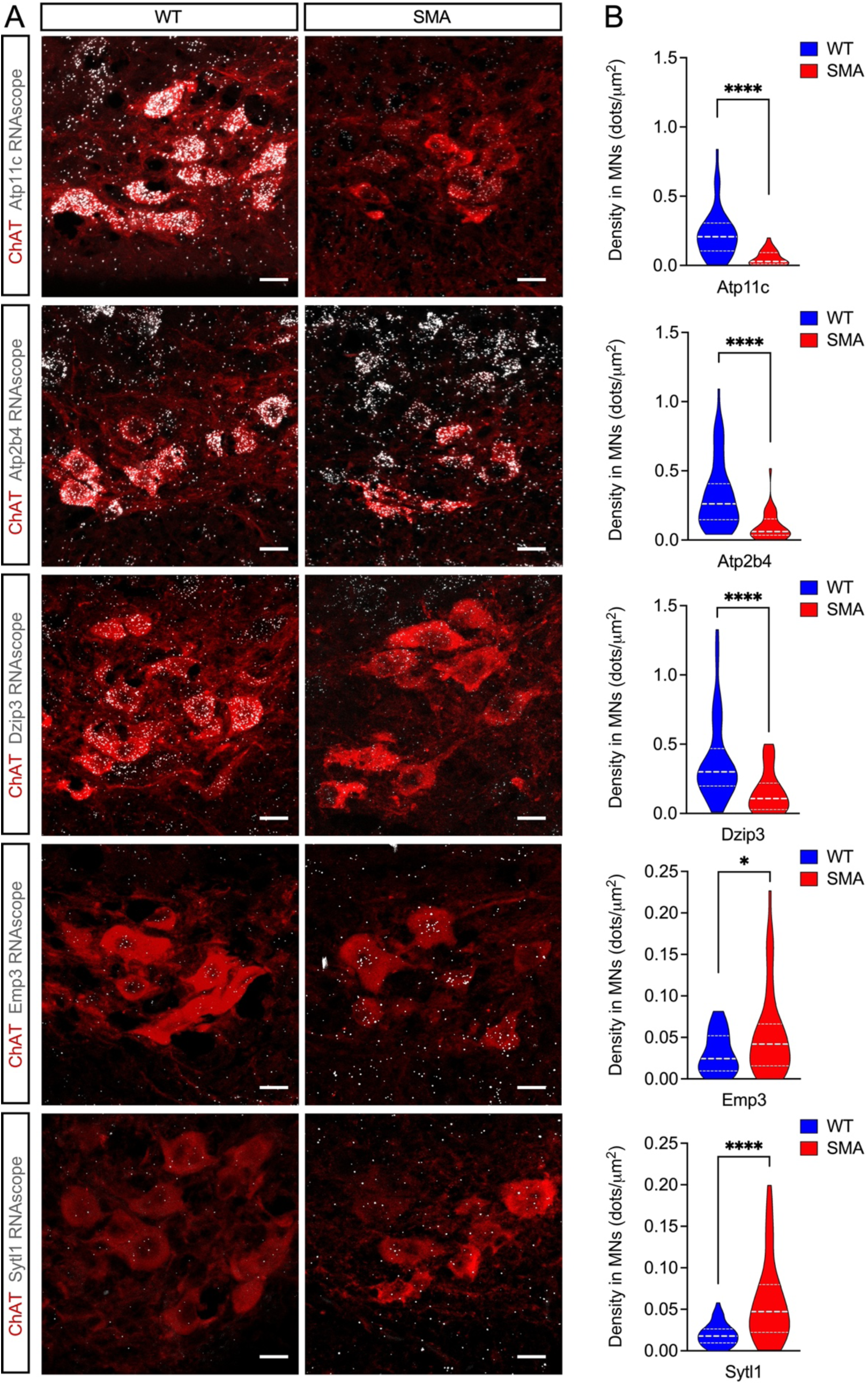
In situ mRNA expression analysis of candidate effectors of p53-dependent neurodegeneration. (**A**) Combined ChAT immunostaining and mRNA detection of the indicated genes by RNAscope in L1 spinal cord sections from WT and SMA mice at P6. Scale bars = 20 µm. (**B**) Violin plots show the number of mRNA foci normalized to the soma area (dots/μm^2^) for each indicated gene from WT (n≥40) and SMA (n≥30) motor neurons. Mann-Whitney test. ****P < 0.0001; *P < 0.05.

**Supplementary Figure S2.**
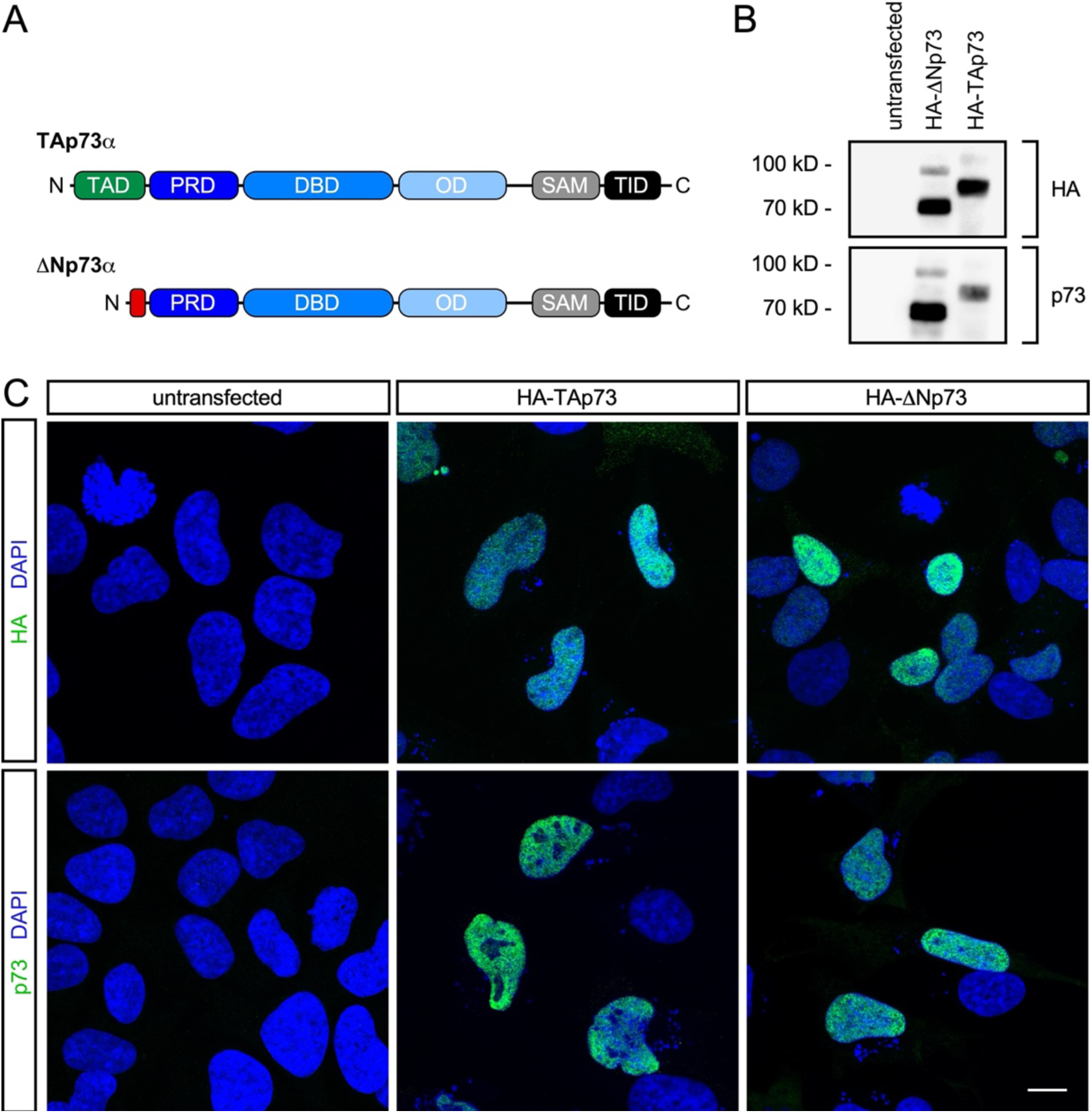
Characterization of the antibody against mouse p73. (**A**) Schematic of the domain structure for the alpha isoforms of TAp73 and ΔNp73 proteins. TAD, transactivation domain; PRD, proline-rich domain; DBD, DNA-binding domain, OD, oligomerization domain, SAM, sterile alpha motif; TID, transcription inhibitory domain. (**B**) Western blot analysis with anti-HA and anti-p73 antibodies of lysates from HEK293 cells that were either untransfected or transfected with plasmids encoding HA-tagged mouse TAp73 or ΔNp73 proteins. (**C**) HeLa cells were either untransfected or transfected with plasmids encoding HA-tagged mouse TAp73 or ΔNp73 proteins followed by immunostaining with anti-HA or anti-p73 antibodies. DAPI was used for nuclear staining. Scale bar = 10 µm.

**Supplementary Figure S3.**
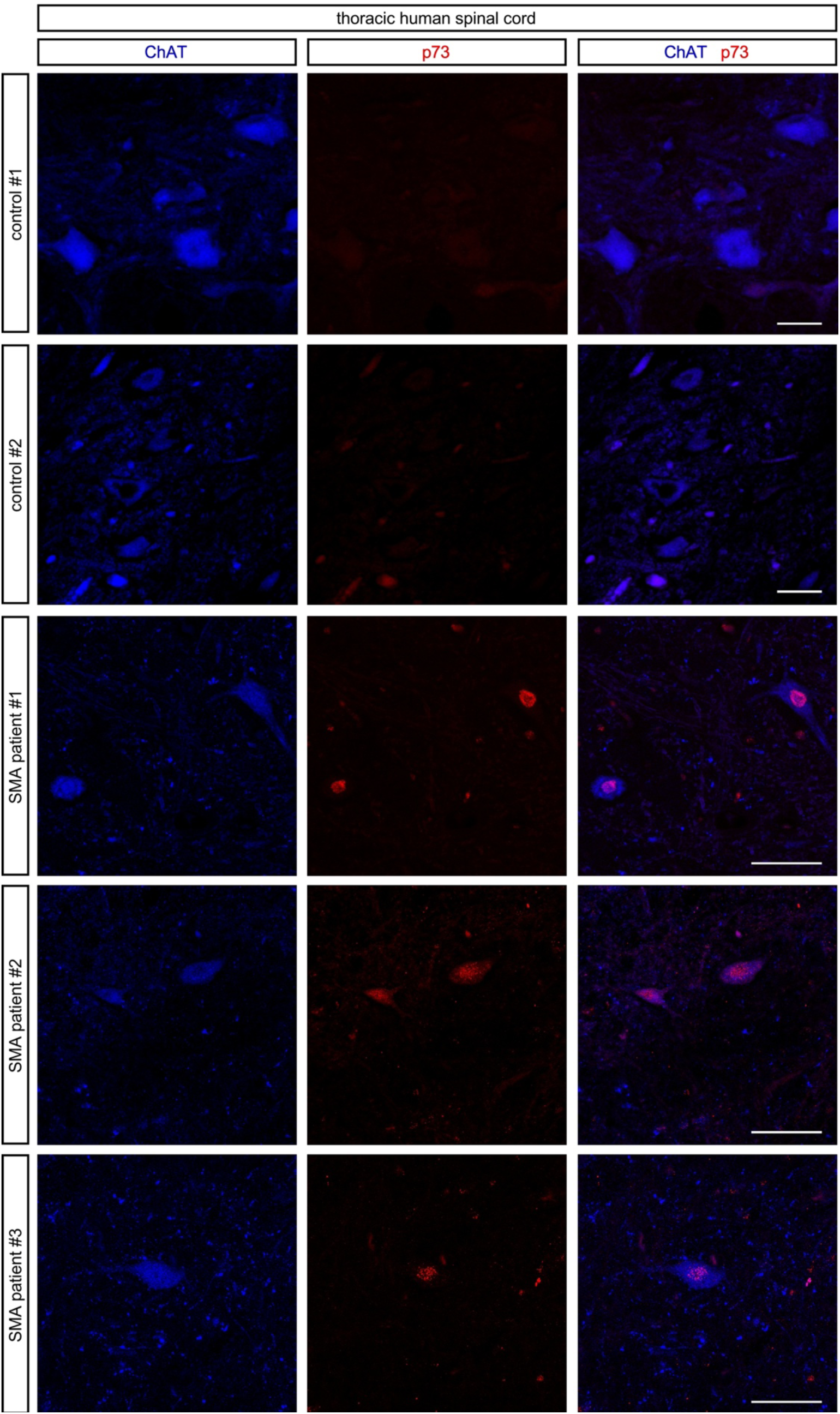
Nuclear accumulation of p73 in motor neurons from type 1 SMA patients. ChAT and p73 immunostaining of thoracic spinal cords isolated postmortem from control and SMA type 1 individuals. Scale bars = 50 µm. See Supplementary Table S2 for further information about the human samples.

**Supplementary Figure S4.**
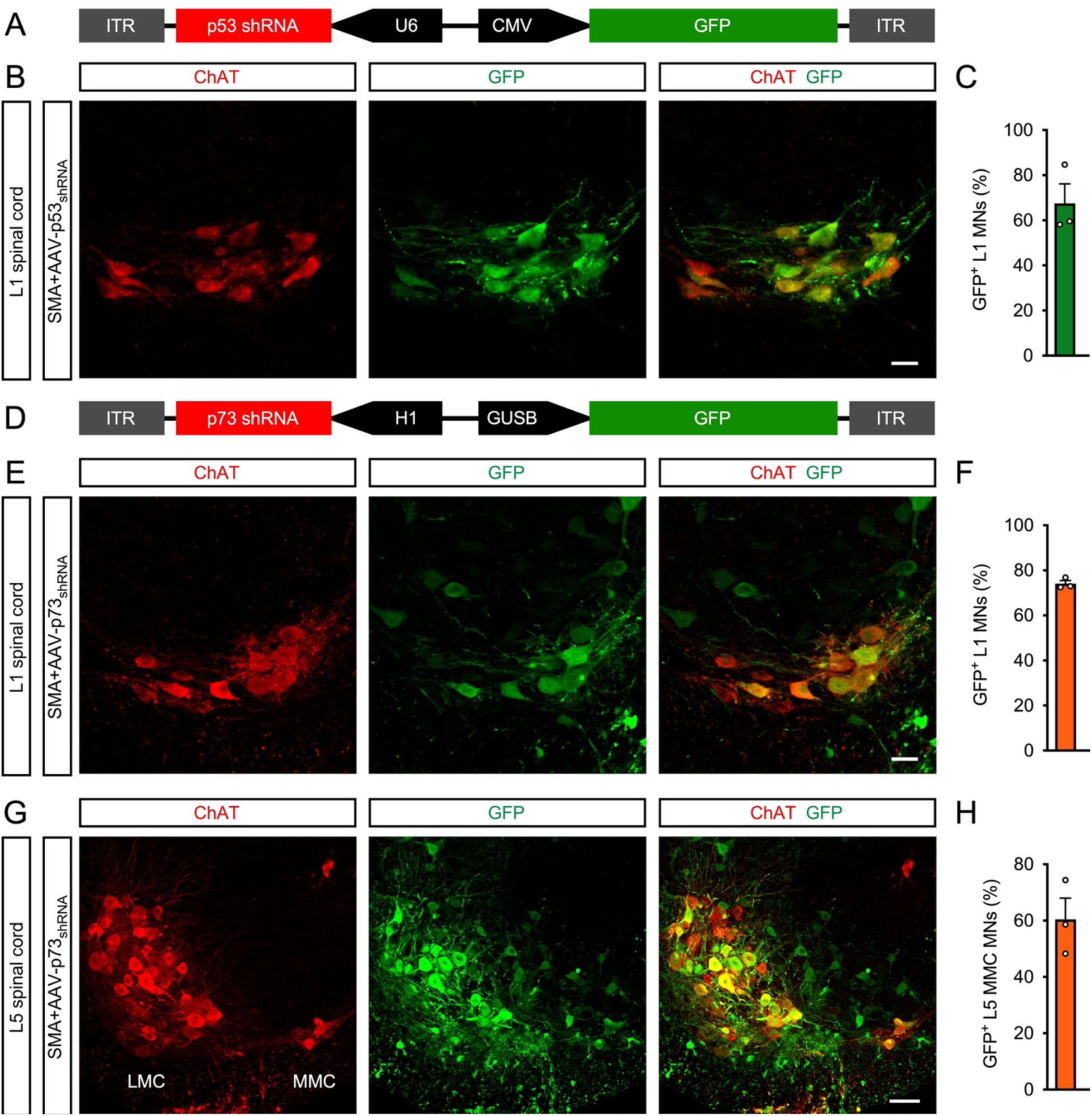
Efficiency of motor neuron transduction by AAV9 vectors in the spinal cord of SMA mice. (**A**) Schematic of the AAV9-p53_shRNA_ vector. (**B**) ChAT and GFP immunostaining of L1 spinal cords isolated at P11 from SMA mice injected with AAV9-p53_shRNA_ at P0. Scale bar = 25 µm. (**C**) Percentage of GFP^+^ L1 motor neurons from experiments as in (B). The graph shows mean, SEM, and individual values from 3 mice. (**D**) Schematic of the AAV9-p73_shRNA_ vector. (**E**) ChAT and GFP immunostaining of L1 spinal cords isolated at P11 from SMA mice injected with AAV9-p73_shRNA_ at P0. Scale bar = 25 µm. (**F**) Percentage of GFP^+^ L1 motor neurons from experiments as in (E). The graph shows mean, SEM, and individual values from 3 mice. (**G**) ChAT and GFP immunostaining of L5 spinal cords isolated at P11 from SMA mice injected with AAV9-p73_shRNA_ at P0. L5 LMC and MMC motor neuron pools are indicated. Scale bar = 50 µm. (**H**) Percentage of GFP^+^ L5 MMC motor neurons from experiments as in (G). The graph shows mean, SEM, and individual values from 3 mice.

**Supplementary Figure S5.**
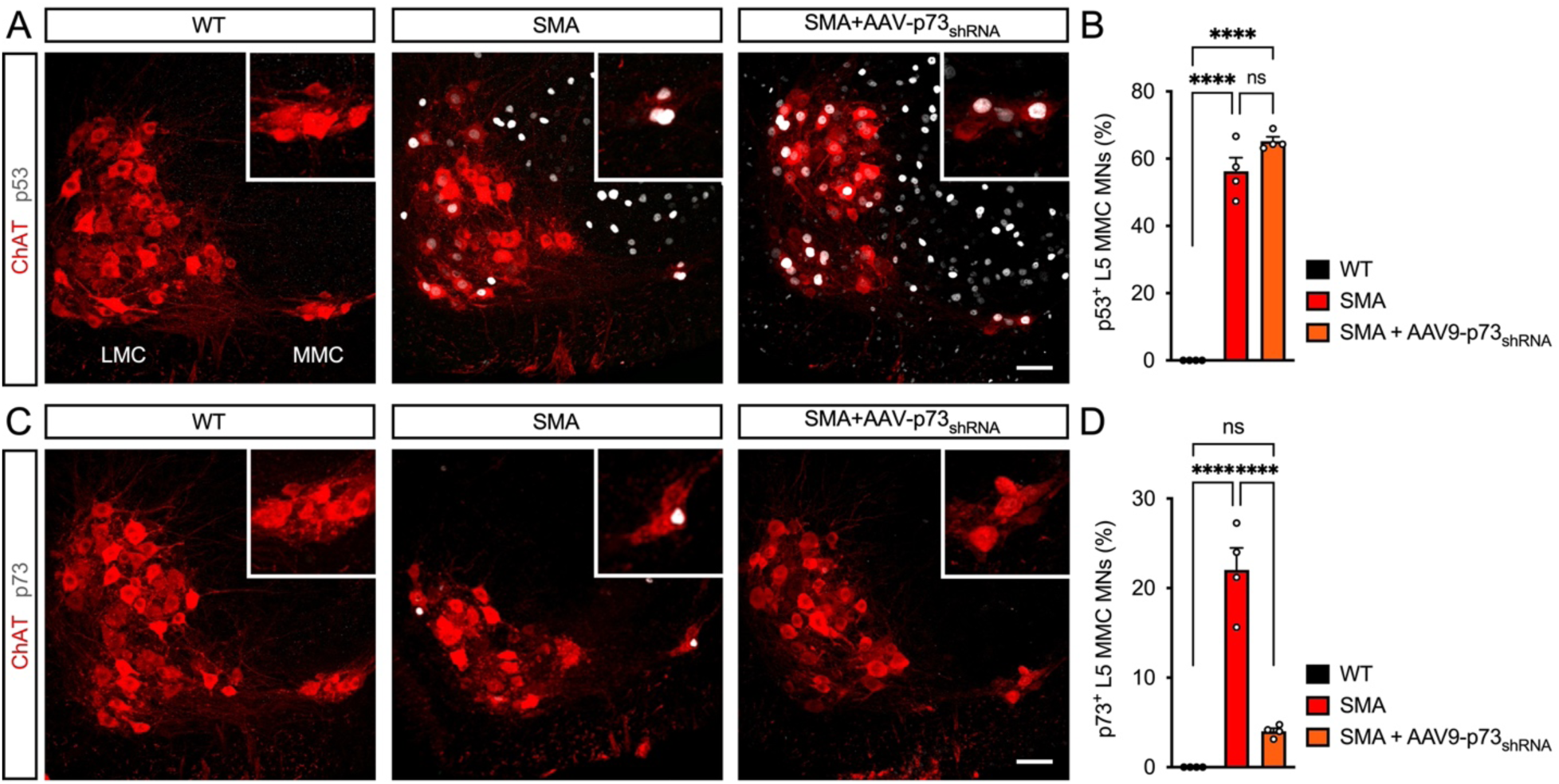
AAV9-mediated p73 knockdown does not affect p53 induction in motor neurons of SMA mice. (**A**) ChAT and p53 immunostaining of L5 spinal cords isolated at P11 from uninjected WT and SMA mice, and SMA mice injected with AAV9-p73_shRNA_ at P0. L5 LMC and MMC motor neuron pools are indicated, and magnified views of L5 MMC motor neurons are shown in the insets. Scale bar = 50 µm. (**B**) Percentage of p53^+^ L5 MMC motor neurons from experiments as in (A). The graph shows mean, SEM, and individual values from WT (n=4), SMA (n=4), and SMA+AAV9-p73_shRNA_ (n=4) mice. One-way ANOVA with Tukey’s *post hoc* test. ****P < 0.0001; ns, not significant. (**C**) ChAT and p73 immunostaining of L5 spinal cords isolated at P11 from uninjected WT and SMA mice, and SMA mice injected with AAV9-p73_shRNA_ at P0. Magnified views of L5 MMC motor neurons are shown in the insets. Scale bar = 50 µm. (**D**) Percentage of p73^+^ L5 MMC motor neurons in the same experimental groups as in (C). The graph shows mean, SEM, and individual values from WT (n=4), SMA (n=4), and SMA+AAV9-p73_shRNA_ (n=4) mice. One-way ANOVA with Tukey’s *post hoc* test. ****P < 0.0001; ns, not significant.

**Supplementary Table S2.** List of human tissue used for immunohistochemical analysis.

| Sample Name | Patient ID | SMA Type | Age | PMI | Analyzed Region |
| --- | --- | --- | --- | --- | --- |
| Control #1 | CNTL_12_06 | Control | 3 years | ~2 h | lumbar, thoracic |
| Control #2 | CNTL_12_04 | Control | 5 years | 21 h | thoracic |
| SMA #1 | SMA_10_01 | Type 1 | 14.8 months | <2 h | lumbar, thoracic |
| SMA #2 | SMA_17_04 | Type 1 | 24 months | 2 h | thoracic |
| SMA #3 | SMA_24_01 | Type 1 | 7 months | 2 h | thoracic |

**Supplementary Table S3.**
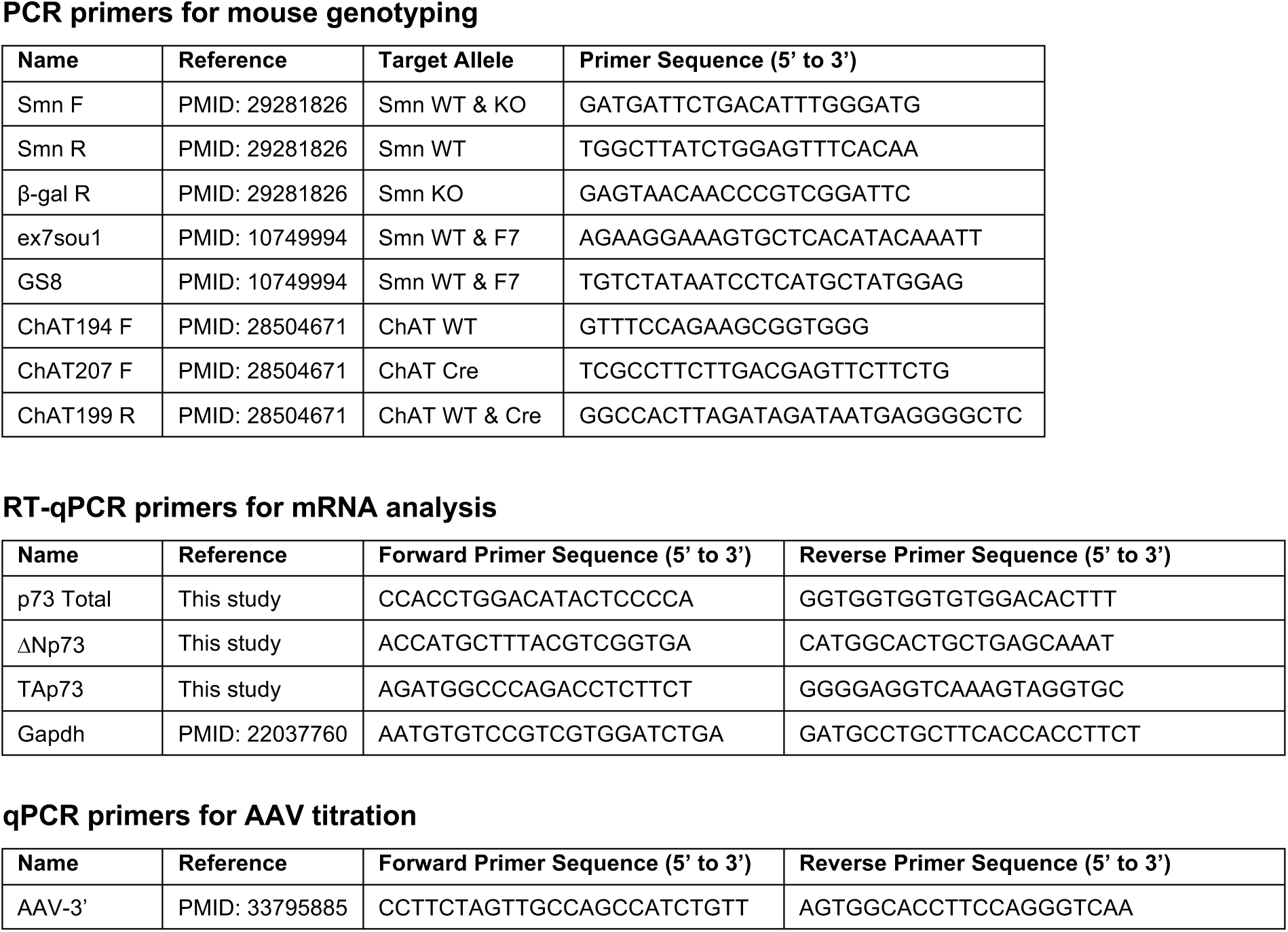
List of primers used in this study.

**Supplementary Table S4.**
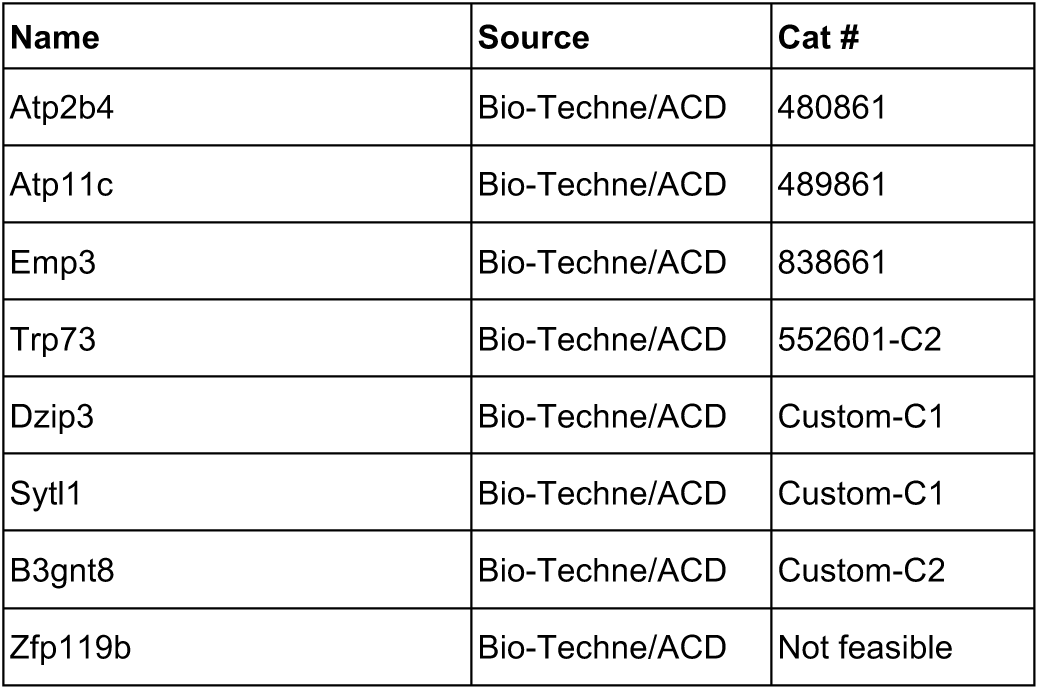
RNAScope probes.

**Supplementary Table S5.** List of antibodies used in this study.

| Name | Source | Cat # | Host | Assay | Dilution |
| --- | --- | --- | --- | --- | --- |
| SMN Clone 8 | BD Transduction Laboratories | 610646 | Mouse | WB | 1:5,000 |
| GFP | Sigma-Aldrich | G1544 | Rabbit | WB | 1:5,000 |
| GAPDH | Sigma-Aldrich | MAB374 | Mouse | WB | 1:50,000 |
| Histone H3 | abcam | ab1791 | Rabbit | WB | 1:50,000 |
| HA-Tag (C29F4) | Cell Signaling | 3724 | Rabbit | WB / ICC | 1:1,000 / 1:1,000 |
| TAp73 (5B429) | Novus Biologicals | NBP2-24737SS | Mouse | WB / ICC | 1:500 / 1:100 |
| p73 delta (38C674.2) | Invitrogen | MA5-16183 | Mouse | WB / ICC | 1:1,000 / 1:100 |
| p73 (ST05-76) | Invitrogen | MA5-32187 | Rabbit | WB / ICC / IHC | 1:1,000 / 1:100 / 1:1000 |
| p53 (CM5) | Leica Biosystems | NCL-p53-CM5p | Rabbit | IHC | 1:1,000 |
| p53 (DO-7) | Cell Signaling | 48818 | Mouse | IHC | 1:250 |
| ChAT | Sigma-Aldrich | AB144P | Goat | IHC | 1:100 |
| GFP | Aves Labs | GFP-1010 | Chicken | IHC | 1:1,000 |
| Synaptophysin1 | Synaptic Systems | 101 308 | Guinea Pig | IHC | 1:500 |
| Neurofilament M | Sigma-Aldrich | AB1987 | Rabbit | IHC | 1:500 |
| Peroxidase AffiniPure™<br>Goat Anti-Mouse | Jackson | 115-035-044 | Goat | WB | 1:10,000 |
| Peroxidase AffiniPure™<br>Goat Anti-Rabbit | Jackson | 111-035-003 | Goat | WB | 1:10,000 |
| Alexa Fluor® 488 AffiniPure®<br>Donkey Anti-Chicken | Jackson | 703-545-155 | Donkey | IHC | 1:250 |
| Alexa Fluor® 488 AffiniPure®<br>Donkey Anti-Rabbit | Jackson | 711-545-152 | Donkey | ICC / IHC | 1:250 / 1:250 |
| Alexa Fluor® 488 AffiniPure®<br>Donkey Anti-Goat | Jackson | 705-545-147 | Donkey | IHC | 1:1,000 |
| Cy™3 AffiniPure®<br>Donkey Anti-Mouse | Jackson | 715-165-150 | Donkey | ICC / IHC | 1:250 / 1:1,000 |
| Cy™3 AffiniPure®<br>Donkey Anti-Rabbit | Jackson | 711-165-152 | Donkey | IHC | 1:250 |
| Cy™5 AffiniPure®<br>Donkey Anti-Goat | Jackson | 705-175-147 | Donkey | IHC | 1:250 |
| Cy™5 AffiniPure®<br>Donkey Anti-Guinea Pig | Jackson | 706-175-148 | Donkey | IHC | 1:250 |
| Alexa Fluor® 647 AffiniPure®<br>Donkey Anti-Rabbit | Jackson | 711-605-152 | Donkey | IHC | 1:1,000 |

## Notes

### Competing Interest Statement

The authors have declared no competing interest.

## References

1. Fu, H., Hardy, J., and Duff, K.E. (2018). Selective vulnerability in neurodegenerative diseases. Nat. Neurosci. 21, 1350–1358. 10.1038/s41593-018-0221-2.

2. Kampmann, M. (2024). Molecular and cellular mechanisms of selective vulnerability in neurodegenerative diseases. Nat. Rev. Neurosci. 25, 351–371. 10.1038/s41583-024-00806-0.

3. Tisdale, S., and Pellizzoni, L. (2015). Disease Mechanisms and Therapeutic Approaches in Spinal Muscular Atrophy. J Neurosci 35, 8691–8700. 10.1523/JNEUROSCI.0417-15.2015.

4. Wirth, B., Karakaya, M., Kye, M.J., and Mendoza-Ferreira, N. (2020). Twenty-Five Years of Spinal Muscular Atrophy Research: From Phenotype to Genotype to Therapy, and What Comes Next. Annu. Rev. Genomics Hum. Genet. 21, 231–261. 10.1146/annurev-genom-102319-103602.

5. Lefebvre, S., Burglen, L., Reboullet, S., Clermont, O., Burlet, P., Viollet, L., Benichou, B., Cruaud, C., Millasseau, P., Zeviani, M., et al. (1995). Identification and characterization of a spinal muscular atrophy-determining gene. Cell 80, 155–165.

6. Lorson, C.L., Hahnen, E., Androphy, E.J., and Wirth, B. (1999). A single nucleotide in the SMN gene regulates splicing and is responsible for spinal muscular atrophy. Proc Natl Acad Sci U A 96, 6307–6311.

7. Lefebvre, S., Burlet, P., Liu, Q., Bertrandy, S., Clermont, O., Munnich, A., Dreyfuss, G., and Melki, J. (1997). Correlation between severity and SMN protein level in spinal muscular atrophy. Nat Genet 16, 265–269.

8. Li, D.K., Tisdale, S., Lotti, F., and Pellizzoni, L. (2014). SMN control of RNP assembly: from post-transcriptional gene regulation to motor neuron disease. Semin Cell Dev Biol 32, 22–29. 10.1016/j.semcdb.2014.04.026.

9. Singh, R.N., Howell, M.D., Ottesen, E.W., and Singh, N.N. (2017). Diverse role of survival motor neuron protein. Biochim. Biophys. Acta - Gene Regul. Mech. 1860, 299–315. 10.1016/j.bbagrm.2016.12.008.

10. Kong, L., Valdivia, D.O., Simon, C.M., Hassinan, C.W., Delestrée, N., Ramos, D.M., Park, J.H., Pilato, C.M., Xu, X., Crowder, M., et al. (2021). Impaired prenatal motor axon development necessitates early therapeutic intervention in severe SMA. Sci. Transl. Med. 13, eabb6871. 10.1126/scitranslmed.abb6871.

11. Lee, J.C., Chung, W.K., Pisapia, D.J., and Henderson, C.E. (2025). Motor pool selectivity of neuromuscular degeneration in type I spinal muscular atrophy is conserved between human and mouse. Hum. Mol. Genet. 34, 347–367. 10.1093/hmg/ddae190.

12. Mentis, G.Z., Blivis, D., Liu, W., Drobac, E., Crowder, M.E., Kong, L., Alvarez, F.J., Sumner, C.J., and O’Donovan, M.J. (2011). Early functional impairment of sensory-motor connectivity in a mouse model of spinal muscular atrophy. Neuron 69, 453–467. 10.1016/j.neuron.2010.12.032.

13. Simon, C.M., Dai, Y., Van Alstyne, M., Koutsioumpa, C., Pagiazitis, J.G., Chalif, J.I., Wang, X., Rabinowitz, J.E., Henderson, C.E., Pellizzoni, L., et al. (2017). Converging Mechanisms of p53 Activation Drive Motor Neuron Degeneration in Spinal Muscular Atrophy. Cell Rep. 21, 3767–3780. 10.1016/j.celrep.2017.12.003.

14. Liu, Y., Su, Z., Tavana, O., and Gu, W. (2024). Understanding the complexity of p53 in a new era of tumor suppression. Cancer Cell 42, 946–967. 10.1016/j.ccell.2024.04.009.

15. Xu, F., Liu, Z., Wang, Z., Zhang, Y., Zhao, Y., and Fang, W. (2025). Progress of p53 Involvement in Central Nervous System Diseases and Targeted Drug Discovery. Cell Biochem. Funct. 43, e70089. 10.1002/cbf.70089.

16. Martin, L.J. (2000). p53 is abnormally elevated and active in the CNS of patients with amyotrophic lateral sclerosis. Neurobiol. Dis. 7, 613–622. 10.1006/nbdi.2000.0314.

17. Ranganathan, S., and Bowser, R. (2010). p53 and Cell Cycle Proteins Participate in Spinal Motor Neuron Cell Death in ALS. Open Pathol. J. 4, 11–22. 10.2174/1874375701004010011.

18. Maor-Nof, M., Shipony, Z., Lopez-Gonzalez, R., Nakayama, L., Zhang, Y.J., Couthouis, J., Blum, J.A., Castruita, P.A., Linares, G.R., Ruan, K., et al. (2021). p53 is a central regulator driving neurodegeneration caused by C9orf72 poly(PR). Cell 184, 689–708.e20. 10.1016/j.cell.2020.12.025.

19. Ziff, O.J., Neeves, J., Mitchell, J., Tyzack, G., Martinez-Ruiz, C., Luisier, R., Chakrabarti, A.M., McGranahan, N., Litchfield, K., Boulton, S.J., et al. (2023). Integrated transcriptome landscape of ALS identifies genome instability linked to TDP-43 pathology. Nat. Commun. 14, 2176. 10.1038/s41467-023-37630-6.

20. Buettner, J.M., Sime Longang, J.K., Gerstner, F., Apel, K.S., Blanco-Redondo, B., Sowoidnich, L., Janzen, E., Langenhan, T., Wirth, B., and Simon, C.M. (2021). Central synaptopathy is the most conserved feature of motor circuit pathology across spinal muscular atrophy mouse models. iScience 24, 103376. 10.1016/j.isci.2021.103376.

21. Carlini, M.J., Triplett, M.K., and Pellizzoni, L. (2022). Neuromuscular denervation and deafferentation but not motor neuron death are disease features in the Smn2B/- mouse model of SMA. PloS One 17. 10.1371/JOURNAL.PONE.0267990.

22. Van Alstyne, M., Simon, C.M., Sardi, S.P., Shihabuddin, L.S., Mentis, G.Z., and Pellizzoni, L. (2018). Dysregulation of Mdm2 and Mdm4 alternative splicing underlies motor neuron death in spinal muscular atrophy. Genes Dev. 32, 1045–1059. 10.1101/gad.316059.118.

23. Simon, C.M., Van Alstyne, M., Lotti, F., Bianchetti, E., Tisdale, S., Watterson, D.M., Mentis, G.Z., and Pellizzoni, L. (2019). Stasimon Contributes to the Loss of Sensory Synapses and Motor Neuron Death in a Mouse Model of Spinal Muscular Atrophy. Cell Rep. 29, 3885–3901.e5. 10.1016/j.celrep.2019.11.058.

24. Carlini, M.J., Espinoza-Derout, J., Van Alstyne, M., Tisdale, S., Workman, E., Triplett, M.K., Tattoli, I., Yadav, S., Henderson, C.E., Watterson, D.M., et al. (2025). Identification of p38 MAPK inhibition as a neuroprotective strategy for combinatorial SMA therapy. EMBO Mol. Med. 17, 2762–2786. 10.1038/s44321-025-00303-6.

25. Mendell, J.R., Al-Zaidy, S., Shell, R., Arnold, W.D., Rodino-Klapac, L.R., Prior, T.W., Lowes, L., Alfano, L., Berry, K., Church, K., et al. (2017). Single-Dose Gene-Replacement Therapy for Spinal Muscular Atrophy. N Engl J Med 377, 1713–1722. 10.1056/NEJMoa1706198.

26. Finkel, R.S., Mercuri, E., Darras, B.T., Connolly, A.M., Kuntz, N.L., Kirschner, J., Chiriboga, C.A., Saito, K., Servais, L., Tizzano, E., et al. (2017). Nusinersen versus Sham Control in Infantile-Onset Spinal Muscular Atrophy. N Engl J Med 377, 1723–1732. 10.1056/NEJMoa1702752.

27. Mercuri, E., Darras, B.T., Chiriboga, C.A., Day, J.W., Campbell, C., Connolly, A.M., Iannaccone, S.T., Kirschner, J., Kuntz, N.L., Saito, K., et al. (2018). Nusinersen versus Sham Control in Later-Onset Spinal Muscular Atrophy. N Engl J Med 378, 625–635. 10.1056/NEJMoa1710504.

28. Baranello, G., Darras, B.T., Day, J.W., Deconinck, N., Klein, A., Masson, R., Mercuri, E., Rose, K., El-Khairi, M., Gerber, M., et al. (2021). Risdiplam in Type 1 Spinal Muscular Atrophy. N. Engl. J. Med. 384, 915–923. 10.1056/nejmoa2009965.

29. Darras, B.T., Masson, R., Mazurkiewicz-Be³dziñska, M., Rose, K., Xiong, H., Zanoteli, E., Baranello, G., Bruno, C., Vlodavets, D., Wang, Y., et al. (2021). Risdiplam-Treated Infants with Type 1 Spinal Muscular Atrophy versus Historical Controls. N. Engl. J. Med. 385, 427–435. 10.1056/nejmoa2102047.

30. Mercuri, E., Pera, M.C., Scoto, M., Finkel, R., and Muntoni, F. (2020). Spinal muscular atrophy — insights and challenges in the treatment era. Nat. Rev. Neurol. 16, 706–715. 10.1038/s41582-020-00413-4.

31. Chaytow, H., Faller, K.M.E., Huang, Y.T., and Gillingwater, T.H. (2021). Spinal muscular atrophy: From approved therapies to future therapeutic targets for personalized medicine. Cell Rep. Med. 2. 10.1016/j.xcrm.2021.100346.

32. Ravi, B., Chan-Cortés, M.H., and Sumner, C.J. (2021). Gene-Targeting Therapeutics for Neurological Disease: Lessons Learned from Spinal Muscular Atrophy. Annu. Rev. Med. 72, 1–14. 10.1146/annurev-med-070119-115459.

33. Zaika, A.I., Slade, N., Erster, S.H., Sansome, C., Joseph, T.W., Pearl, M., Chalas, E., and Moll, U.M. (2002). DeltaNp73, a dominant-negative inhibitor of wild-type p53 and TAp73, is up-regulated in human tumors. J. Exp. Med. 196, 765–780. 10.1084/jem.20020179.

34. Rufini, A., Agostini, M., Grespi, F., Tomasini, R., Sayan, B.S., Niklison-Chirou, M.V., Conforti, F., Velletri, T., Mastino, A., Mak, T.W., et al. (2011). p73 in Cancer. Genes Cancer 2, 491–502. 10.1177/1947601911408890.

35. Naryshkin, N.A., Weetall, M., Dakka, A., Narasimhan, J., Zhao, X., Feng, Z., Ling, K.K., Karp, G.M., Qi, H., Woll, M.G., et al. (2014). Motor neuron disease. SMN2 splicing modifiers improve motor function and longevity in mice with spinal muscular atrophy. Science 345, 688–693. 10.1126/science.1250127.

36. Lotti, F., Imlach, W.L., Saieva, L., Beck, E.S., le, T.H., Li, D.K., Jiao, W., Mentis, G.Z., Beattie, C.E., McCabe, B.D., et al. (2012). An SMN-Dependent U12 Splicing Event Essential for Motor Circuit Function. Cell 151, 440–454. 10.1016/j.cell.2012.09.012.

37. Ottesen, E.W., Singh, N.N., Luo, D., Kaas, B., Gillette, B.J., Seo, J., Jorgensen, H.J., and Singh, R.N. (2023). Diverse targets of SMN2-directed splicing-modulating small molecule therapeutics for spinal muscular atrophy. Nucleic Acids Res. 51, 5948–5980. 10.1093/nar/gkad259.

38. Nemajerova, A., and Moll, U.M. (2019). Tissue-specific roles of p73 in development and homeostasis. J. Cell Sci. 132, jcs233338. 10.1242/jcs.233338.

39. Logotheti, S., Richter, C., Murr, N., Spitschak, A., Marquardt, S., and Pützer, B.M. (2021). Mechanisms of Functional Pleiotropy of p73 in Cancer and Beyond. Front. Cell Dev. Biol. 9, 737735. 10.3389/fcell.2021.737735.

40. Vikhreva, P., Melino, G., and Amelio, I. (2018). p73 Alternative Splicing: Exploring a Biological Role for the C-Terminal Isoforms. J. Mol. Biol. 430, 1829–1838. 10.1016/j.jmb.2018.04.034.

41. Fuertes-Alvarez, S., Maeso-Alonso, L., Villoch-Fernandez, J., Wildung, M., Martin-Lopez, M., Marshall, C., Villena-Cortes, A.J., Diez-Prieto, I., Pietenpol, J.A., Tissir, F., et al. (2018). p73 regulates ependymal planar cell polarity by modulating actin and microtubule cytoskeleton. Cell Death Dis. 9, 1183. 10.1038/s41419-018-1205-6.

42. Fujitani, M., Sato, R., and Yamashita, T. (2017). Loss of p73 in ependymal cells during the perinatal period leads to aqueductal stenosis. Sci. Rep. 7, 12007. 10.1038/s41598-017-12105-z.

43. Frugier, T., Tiziano, F.D., Cifuentes-Diaz, C., Miniou, P., Roblot, N., Dierich, A., Meur, M.L., and Melki, J. (2000). Nuclear targeting defect of SMN lacking the C-terminus in a mouse model of spinal muscular atrophy. Hum Mol Genet 9, 849–858.

44. Martinez, T.L., Kong, L., Wang, X., Osborne, M.A., Crowder, M.E., Meerbeke, J.P.V., Xu, X., Davis, C., Wooley, J., Goldhamer, D.J., et al. (2012). Survival motor neuron protein in motor neurons determines synaptic integrity in spinal muscular atrophy. J Neurosci 32, 8703–8715. 10.1523/jneurosci.0204-12.2012.

45. Fletcher, E.V., Simon, C.M., Pagiazitis, J.G., Chalif, J.I., Vukojicic, A., Drobac, E., Wang, X., and Mentis, G.Z. (2017). Reduced sensory synaptic excitation impairs motor neuron function via Kv2.1 in spinal muscular atrophy. Nat Neurosci 20, 905–916. 10.1038/nn.4561.

46. Tisdale, S., Van Alstyne, M., Simon, C.M., Mentis, G.Z., and Pellizzoni, L. (2022). SMN controls neuromuscular junction integrity through U7 snRNP. Cell Rep. 40, 111393. 10.1016/J.CELREP.2022.111393.

47. He, H., Wang, C., Dai, Q., Li, F., Bergholz, J., Li, Z., Li, Q., and Xiao, Z.-X. (2016). p53 and p73 Regulate Apoptosis but Not Cell-Cycle Progression in Mouse Embryonic Stem Cells upon DNA Damage and Differentiation. Stem Cell Rep. 7, 1087–1098. 10.1016/j.stemcr.2016.10.008.

48. Pozniak, C.D., Barnabé-Heider, F., Rymar, V.V., Lee, A.F., Sadikot, A.F., and Miller, F.D. (2002). p73 is required for survival and maintenance of CNS neurons. J. Neurosci. Off. J. Soc. Neurosci. 22, 9800–9809. 10.1523/JNEUROSCI.22-22-09800.2002.

49. Tissir, F., Ravni, A., Achouri, Y., Riethmacher, D., Meyer, G., and Goffinet, A.M. (2009). DeltaNp73 regulates neuronal survival in vivo. Proc. Natl. Acad. Sci. U. S. A. 106, 16871–16876. 10.1073/pnas.0903191106.

50. Kartasheva, N.N., Lenz-Bauer, C., Hartmann, O., Schäfer, H., Eilers, M., and Dobbelstein, M. (2003). DeltaNp73 can modulate the expression of various genes in a p53-independent fashion. Oncogene 22, 8246–8254. 10.1038/sj.onc.1207138.

51. Alonso-Olivares, H., Marques, M.M., Prieto-Colomina, A., López-Ferreras, L., Martínez-García, N., Vázquez-Jiménez, A., Borrell, V., Marin, M.C., and Fernandez-Alonso, R. (2024). Mouse cortical organoids reveal key functions of p73 isoforms: TAp73 governs the establishment of the archetypical ventricular-like zones while DNp73 is central in the regulation of neural cell fate. Front. Cell Dev. Biol. 12, 1464932. 10.3389/fcell.2024.1464932.

52. Möller, S., Spitschak, A., Murr, N., and Pützer, B.M. (2025). Integrative Multimodal Profiling of TAp73 and DNp73 Reveals Isoform-Specific Transcriptomic Coregulator Landscapes in Cancer Programs. Biomolecules 16, 63. 10.3390/biom16010063.

53. Sanchez-Prieto, R., Sanchez-Arevalo, V.J., Servitja, J.-M., and Gutkind, J.S. (2002). Regulation of p73 by c-Abl through the p38 MAP kinase pathway. Oncogene 21, 974–979. 10.1038/sj.onc.1205134.

54. Indeglia, A., Valdespino, A., Pantella, G., Offley, S., Hill, C., Foster, M., Casey, K., Tang, H.-Y., Faustino, A.M., Gardini, A., et al. (2025). Targeted citrullination enables p53 binding to non-canonical sites. Mol. Cell 85, 3588–3604.e11. 10.1016/j.molcel.2025.09.004.

55. Stafford, N., Wilson, C., Oceandy, D., Neyses, L., and Cartwright, E.J. (2017). The Plasma Membrane Calcium ATPases and Their Role as Major New Players in Human Disease. Physiol. Rev. 97, 1089–1125. 10.1152/physrev.00028.2016.

56. Sakuragi, T., and Nagata, S. (2023). Regulation of phospholipid distribution in the lipid bilayer by flippases and scramblases. Nat. Rev. Mol. Cell Biol. 24, 576–596. 10.1038/s41580-023-00604-z.

57. Emmrich, S., Wang, W., John, K., Li, W., and Pützer, B.M. (2009). Antisense gapmers selectively suppress individual oncogenic p73 splice isoforms and inhibit tumor growth in vivo. Mol. Cancer 8, 61. 10.1186/1476-4598-8-61.

58. Valsecchi, V., Errico, F., Bassareo, V., Marino, C., Nuzzo, T., Brancaccio, P., Laudati, G., Casamassa, A., Grimaldi, M., D’Amico, A., et al. (2023). SMN deficiency perturbs monoamine neurotransmitter metabolism in spinal muscular atrophy. Commun. Biol. 6. 10.1038/s42003-023-05543-1.

59. Pagiazitis, J.G., Delestrée, N., Sowoidnich, L., Sivakumar, N., Simon, C.M., Chatzisotiriou, A., Albani, M., and Mentis, G.Z. (2025). Catecholaminergic dysfunction drives postural and locomotor deficits in a mouse model of spinal muscular atrophy. Cell Rep. 44, 115147. 10.1016/j.celrep.2024.115147.

60. Gerstner, F., Wittig, S., Menedo, C., Ruwald, S., Carlini, M.J., Vankova, A., Sowoidnich, L., Martín-López, G., Dreilich, V., Alonso-Collado, A., et al. (2026). Cerebellar pathology contributes to neurodevelopmental deficits in spinal muscular atrophy. Brain J. Neurol. 149, 840–855. 10.1093/brain/awaf336.

61. Bae, B.-I., Xu, H., Igarashi, S., Fujimuro, M., Agrawal, N., Taya, Y., Hayward, S.D., Moran, T.H., Montell, C., Ross, C.A., et al. (2005). p53 mediates cellular dysfunction and behavioral abnormalities in Huntington’s disease. Neuron 47, 29–41. 10.1016/j.neuron.2005.06.005.

62. Mansky, R.H., Greguske, E.A., Yu, D., Zarate, N., Intihar, T.A., Tsai, W., Brown, T.G., Thayer, M.N., Kumar, K., and Gomez-Pastor, R. (2023). Tumor suppressor p53 regulates heat shock factor 1 protein degradation in Huntington’s disease. Cell Rep. 42, 112198. 10.1016/j.celrep.2023.112198.

63. da Costa, C.A., Sunyach, C., Giaime, E., West, A., Corti, O., Brice, A., Safe, S., Abou-Sleiman, P.M., Wood, N.W., Takahashi, H., et al. (2009). Transcriptional repression of p53 by parkin and impairment by mutations associated with autosomal recessive juvenile Parkinson’s disease. Nat. Cell Biol. 11, 1370–1375. 10.1038/ncb1981.

64. Duplan, E., Giordano, C., Checler, F., and Alves da Costa, C. (2016). Direct á-synuclein promoter transactivation by the tumor suppressor p53. Mol. Neurodegener. 11, 13. 10.1186/s13024-016-0079-2.

65. Qi, X., Davis, B., Chiang, Y.H., Filichia, E., Barnett, A., Greig, N.H., Hoffer, B., and Luo, Y. (2016). Dopaminergic neuron-specific deletion of p53 gene is neuroprotective in an experimental Parkinson’s disease model. J. Neurochem. 138, 746–757. 10.1111/jnc.13706.

66. de la Monte, S.M., Sohn, Y.K., and Wands, J.R. (1997). Correlates of p53- and Fas (CD95)-mediated apoptosis in Alzheimer’s disease. J. Neurol. Sci. 152, 73–83. 10.1016/s0022-510x(97)00131-7.

67. Ohyagi, Y., Asahara, H., Chui, D.-H., Tsuruta, Y., Sakae, N., Miyoshi, K., Yamada, T., Kikuchi, H., Taniwaki, T., Murai, H., et al. (2005). Intracellular Abeta42 activates p53 promoter: a pathway to neurodegeneration in Alzheimer’s disease. FASEB J. Off. Publ. Fed. Am. Soc. Exp. Biol. 19, 255–257. 10.1096/fj.04-2637fje.

68. Wan, L., Yang, F., Yin, A., Luo, Y., Liu, Y., Liu, F., Wang, J.-Z., Liu, R., and Wang, X. (2025). Age-related p53 SUMOylation accelerates senescence and tau pathology in Alzheimer’s disease. Cell Death Differ. 32, 837–854. 10.1038/s41418-025-01448-0.

69. Culmsee, C., Zhu, X., Yu, Q.S., Chan, S.L., Camandola, S., Guo, Z., Greig, N.H., and Mattson, M.P. (2001). A synthetic inhibitor of p53 protects neurons against death induced by ischemic and excitotoxic insults, and amyloid beta-peptide. J. Neurochem. 77, 220–228. 10.1046/j.1471-4159.2001.t01-1-00220.x.

70. Yang, L.-Y., Greig, N.H., Huang, Y.-N., Hsieh, T.-H., Tweedie, D., Yu, Q.-S., Hoffer, B.J., Luo, Y., Kao, Y.-C., and Wang, J.-Y. (2016). Post-traumatic administration of the p53 inactivator pifithrin-α oxygen analogue reduces hippocampal neuronal loss and improves cognitive deficits after experimental traumatic brain injury. Neurobiol. Dis. 96, 216–226. 10.1016/j.nbd.2016.08.012.

71. Le, T.T., Pham, L.T., Butchbach, M.E., Zhang, H.L., Monani, U.R., Coovert, D.D., Gavrilina, T.O., Xing, L., Bassell, G.J., and Burghes, A.H. (2005). SMNDelta7, the major product of the centromeric survival motor neuron (SMN2) gene, extends survival in mice with spinal muscular atrophy and associates with full-length SMN. Hum Mol Genet 14, 845–857.

72. Van Alstyne, M., Tattoli, I., Delestrée, N., Recinos, Y., Workman, E., Shihabuddin, L.S., Zhang, C., Mentis, G.Z., and Pellizzoni, L. (2021). Gain of toxic function by long-term AAV9-mediated SMN overexpression in the sensorimotor circuit. Nat. Neurosci. 24, 930–940. 10.1038/s41593-021-00827-3.

73. Ruggiu, M., McGovern, V.L., Lotti, F., Saieva, L., Li, D.K., Kariya, S., Monani, U.R., Burghes, A.H., and Pellizzoni, L. (2012). A role for SMN exon 7 splicing in the selective vulnerability of motor neurons in spinal muscular atrophy. Mol Cell Biol 32, 126–138. 10.1128/MCB.06077-11.

